# Progenitor-intrinsic Metabolic Sensing Promotes Hematopoietic Homeostasis

**DOI:** 10.1101/2021.09.20.461098

**Authors:** Hannah A. Pizzato, Yahui Wang, Michael J. Wolfgang, Brian N. Finck, Gary J. Patti, Deepta Bhattacharya

## Abstract

Hematopoietic homeostasis is maintained by stem and progenitor cells in part by extrinsic feedback cues triggered by mature cell loss. We demonstrate a different mechanism by which hematopoietic progenitors intrinsically anticipate and prevent the loss of mature progeny through metabolic switches. We examined hematopoiesis in mice conditionally deficient in long-chain fatty acid oxidation (carnitine palmitoyltransferase 2, *Cpt2*), glutaminolysis (glutaminase, *Gls*), or mitochondrial pyruvate import (mitochondrial pyruvate carrier 2, *Mpc2*). While genetic ablation of *Cpt2* or *Gls* minimally impacted most blood lineages, deletion of *Mpc2* led to a sharp decline in mature myeloid cells. However, MPC2-deficient myeloid cells rapidly recovered due to a transient increase in myeloid progenitor proliferation. Competitive bone marrow chimera and stable isotope tracing experiments demonstrated that this proliferative burst was intrinsic to MPC2-deficient progenitors and accompanied by a metabolic switch to glutaminolysis. Thus, hematopoietic progenitors intrinsically adjust to metabolic perturbations independently of feedback from downstream mature cells to maintain homeostasis.

## Introduction

As hematopoietic stem cells (HSCs) differentiate, they gradually lose the potential to generate specific blood cell types until commitment to a single lineage is achieved. Since HSCs are exceedingly rare, their differentiation occurs concomitantly with a progressive expansion of downstream progenitors to ensure sufficient production of mature cells (Bryder et al., 2006). Along with a numerical expansion, downstream progenitors possess both more proliferative and differentiative capacity than do HSCs (Passegué et al., 2005; Sun et al., 2014). This progressive expansion of progenitors and mature lineages is known as transit amplification. Under homeostatic conditions, transit amplification matches the rate of new cell generation with the turnover of mature cells. Because each mature lineage has its own distinct lifespan, the mechanisms that maintain homeostasis are necessarily complicated. For example, neutrophils do not survive for more than a few days, while mature B lymphocytes persist for several months (Cronkite et al., 1959; Fulcher & Basten, 1997; Mauer et al., 1960; Pillay et al., 2010). The different arms of hematopoiesis must therefore be controlled in modular and separable ways.

Feedback triggered by a loss of downstream cells to upstream progenitors is one well-established mechanism by which homeostasis is maintained. As one example, repeated bleeding of mice triggers HSCs to proliferate and self-renew at an increased rate, leading to a restoration of bone marrow cells and increased splenocytes (Cheshier et al., 2007). Upon transfusion of red blood cells (RBCs) in these mice, HSC proliferation returns to normal. As another example, transplantation of a small number of wild-type HSCs into RAG2 and IL-2 receptor common γ chain-deficient mice, which lack B, T, and natural killer (NK) cells, results in a rapid repopulation of mature lymphocytes (Bhattacharya et al., 2006). This is driven by a pronounced and selective expansion of B and T cell progenitors despite very low levels of HSC chimerism. These data suggest that HSCs and progenitors respond to fill a void left by the loss of more differentiated downstream progenitors and mature cells. Yet in these experimental settings, progenitor adaptations were observed only after substantial depletion of their downstream lineages. To maintain homeostasis under normal conditions, it would seem advantageous for progenitors to anticipate and adjust in real time to perturbations that might affect more differentiated progeny. Such a mechanism would not require a dramatic loss of mature lineages before progenitors would adjust their output.

Metabolic adaption might represent one situation where hematopoietic progenitors adjust output in response to perturbations that impact downstream mature cells. For example, a decrease in lymphocytes occurs within hours of streptozotocin treatment (Muller et al., 2011), which ablates pancreatic beta cells and induces diabetes and high glucose levels (Arison et al., 1967; Lenzen, 2008; Rakieten et al., 1963). Yet by 4 weeks after treatment with streptozotocin, lymphocyte numbers are normal (Nagareddy et al., 2013). Similarly, mouse lymphocytes decrease after 6 weeks on high fat diet (Luo et al., 2015) but increase after 90 days (Trottier et al., 2012). Moreover, nutrient availability varies substantially throughout the course of the day (Pickel & Sung, 2020; Potter et al., 2016; Schlierf & Dorow, 1973), yet there is relatively little impact on the abundances of mature cells (Born et al., 1997; Haus et al., 1983; Lawrence et al., 1933; Sabin et al., 1927). Though these examples represent independent studies and are not directly comparable to each other, the data hint that while severe metabolic alterations may transiently alter hematopoietic composition, progenitors may be able to adapt to these changes to recover and maintain homeostasis. Defining whether and how such an adaptation occurs requires genetic tools and a dissection of progenitor-intrinsic metabolic pathways.

Much of the work performed to date on hematopoietic metabolism has focused on the relative dependence on glycolysis versus mitochondrial functions by HSCs and progenitors. The metabolic profile of HSCs is notably different than those of lineage-committed progenitors (Agathocleous et al., 2017). HSCs often rely on glycolysis to remain quiescent, but as differentiation proceeds, a shift to increased oxidative phosphorylation occurs to ensure bioenergetic requirements are met (Takubo et al., 2013; Yu et al., 2012; Wang et al., 2014; Maryanovich et al., 2015; Ito & Suda, 2014; Nakamura-Ishizu et al., 2020; Simsek et al., 2010; Suda et al., 2011; Ho et al., 2017). Accordingly, lineage-committed progenitors produce more reactive oxygen species than do upstream HSCs, and increasing levels of reactive oxygen species hinders HSC function (Inoue et al., 2010; Ito et al., 2004, 2006; Norddahl et al., 2011; Tan & Suda, 2018; Tothova et al., 2007; Wang et al., 2014). Though HSCs do still rely on mitochondrial functions (Ansó et al., 2017; Bejarano-García et al., 2016; Chen et al., 2008; Gan et al., 2010; Guitart et al., 2017; Gurumurthy et al., 2010; Hinge et al., 2020; Inoue et al., 2010; Ito et al., 2019; Luchsinger et al., 2016; Mansell et al., 2021; Maryanovich et al., 2012; Mortensen et al., 2011; Nakada et al., 2010; Norddahl et al., 2011; Qi et al., 2021; Umemoto et al., 2018; Vannini et al., 2016), the preponderance of evidence suggests that hematopoietic progenitors depend more heavily on oxidative phosphorylation of carbon sources to generate ATP. Yet these specific carbon sources have not yet been genetically defined *in vivo*.

Here, we utilized *in vivo* genetic models to interrogate the roles of long-chain fatty acid oxidation, glutaminolysis, and mitochondrial pyruvate import in hematopoietic homeostasis. In a striking example of homeostatic maintenance, myeloid progenitors genetically deprived of mitochondrial pyruvate not only switched to glutaminolysis, but this switch was also accompanied by a transient and rapid proliferative burst to regenerate themselves and their mature myeloid daughters. These data demonstrate that progenitors have intrinsic metabolic sensing abilities that promote hematopoietic homeostasis.

## Results

### Long-chain fatty acid oxidation and glutaminolysis are dispensable for most hematopoietic lineages

To define essential carbon sources that fuel the tricarboxylic acid (TCA) cycle in hematopoietic lineages, we employed *in vivo* genetic ablation models. We focused on long-chain fatty acid oxidation, glutaminolysis, and mitochondrial pyruvate utilization, as each of these pathways are major contributors to ATP production in many other cell types. First, we used mice deficient in carnitine palmitoyltransferase 2 (CPT2), which transports long-chain fatty acids into the inner matrix of the mitochondria for oxidation (Houten et al., 2016; Lee et al., 2015). Because germline deletion of *Cpt2* is lethal (Isackson et al., 2008; Ji et al., 2008; Longo et al., 2006; Nyman et al., 2005), we crossed *Cpt2^fl/fl^* mice (Lee et al., 2015) to *ROSA26 CreER* animals, which constitutively express tamoxifen-inducible Cre recombinase (Ventura et al., 2007). Wild-type CD45.1^+^ and *Cpt2^fl/fl^; ROSA26 CreER^-/-^* or *^+/+^* CD45.2^+^ bone marrow cells were mixed in equal proportions and transplanted into irradiated recipients (**Figure 1A**). Reconstitution was allowed to proceed for at least 8 weeks before administration of tamoxifen to ablate *Cpt2* in CD45.2^+^ cells. This system avoids lethality following *Cpt2* deletion and allows assessment of blood lineage-intrinsic phenotypes.

**Figure 1.**
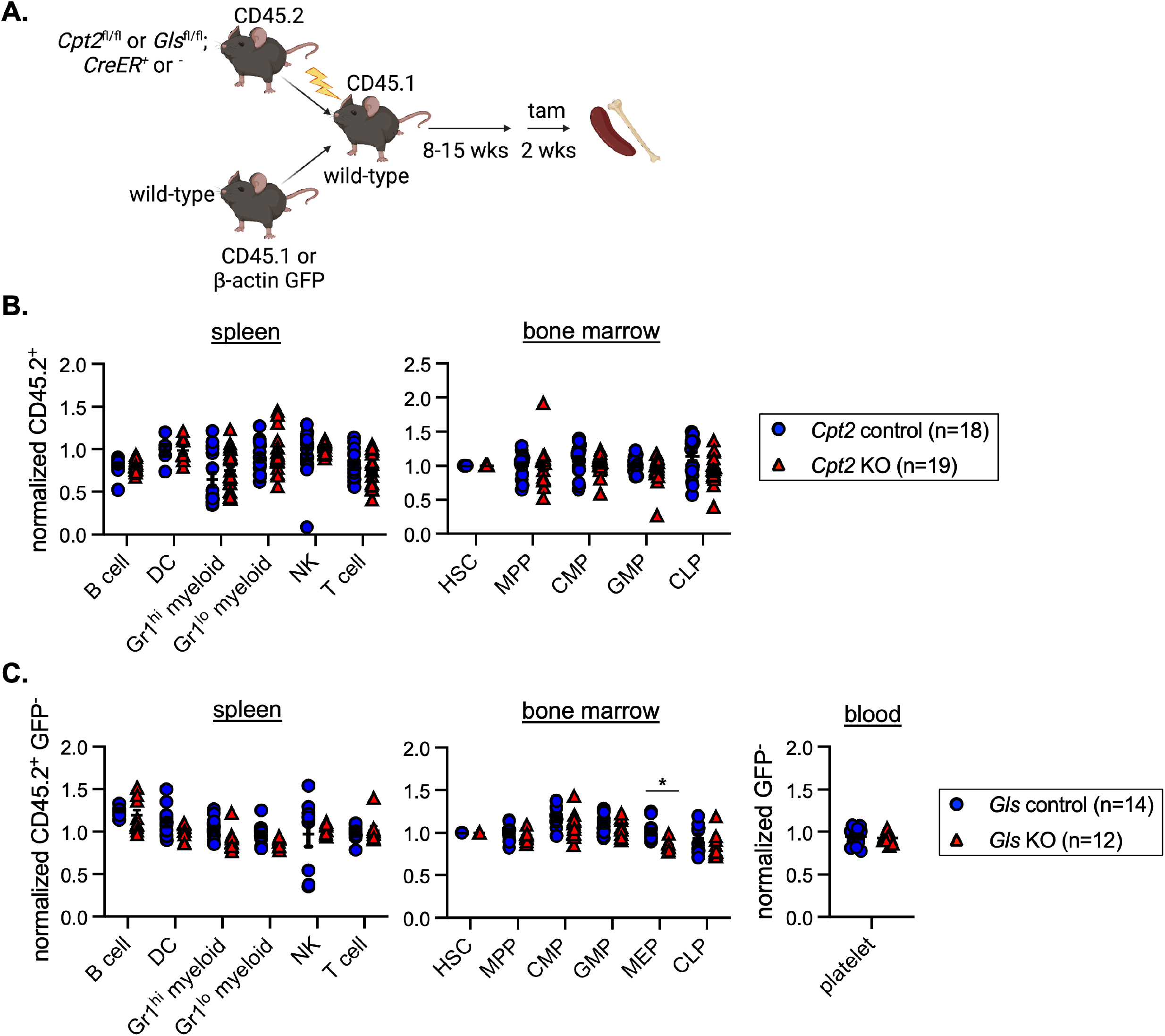
Long-chain fatty acid oxidation and glutaminolysis are dispensable for most hematopoietic lineages. (A) Schematic representation of mixed bone marrow chimera experiments to assess hematopoietic requirements for *Cpt2* control (*Cpt2^fl/fl^; ROSA26 CreER^-/-^*) or KO (*Cpt2^fl/fl^; ROSA26 CreER^+/+^*) cells and *Gls* control (*Gls^fl/fl^; ROSA26 CreER^-/-^*) or KO (*Gls^fl/fl^; ROSA26 CreER^+/+^*) cells. (B) CD45.2 chimerism was normalized to either pre-tamoxifen peripheral blood chimerism levels for the spleen or hematopoietic stem cell (HSC) chimerism for the bone marrow of *Cpt2* chimeras. Data are shown for B cells, dendritic cells (DCs), Gr1^hi^ and ^lo^ myeloid cells, natural killer (NK) cells, and T cells for the spleen. Data are shown for HSCs, multipotent progenitors (MPPs), common myeloid progenitors (CMPs), granulocyte-macrophage progenitors (GMPs), and common lymphoid progenitors (CLPs) in the bone marrow. Each symbol represents an individual mouse. Mean values ± SEM are shown, and data are pooled from three independent experiments. (C) CD45.2^+^ GFP^-^ chimerism was normalized as stated in (B) for *Gls* chimeras. Cell populations assessed are the same as in (B) with the addition of megakaryocyte-erythroid progenitors (MEPs) in the bone marrow and platelets in the blood. Each symbol represents an individual mouse. Mean values ± SEM are shown, and data are pooled from two independent experiments. *p<0.05 by 2-way ANOVA with post-hoc Tukey’s multiple comparisons test.

After 2 weeks of tamoxifen treatment, chimerism of hematopoietic populations in the spleen and bone marrow was assessed and normalized to the chimerism of either pre-tamoxifen blood lineages for splenic populations or hematopoietic stem cells for bone marrow progenitors. Analysis of the spleen included B cells, dendritic cells (DCs), Gr1^hi^ myeloid cells, Gr1^lo^ myeloid cells, natural killer (NK) cells, and T cells (gating strategy in **Figure S1A**). Bone marrow populations included hematopoietic stem cells (HSCs), multipotent progenitors (MPPs), Flk2^+^ common myeloid progenitors (CMPs), granulocyte-macrophage progenitors (GMPs), and common lymphoid progenitors (CLPs) (gating strategy in **Figure S1B**). We found no differences between *Cpt2* knockout cells and wild-type controls (**Figure 1B**), despite efficient *Cpt2* deletion (**Figure S1C**). This suggests that long-chain fatty acid oxidation is not required for the development or maintenance of lymphoid or myeloid cells, and that other pathways such as glutaminolysis may be more important for these lineages. Prior studies have suggested an important role for glutaminolysis in human erythropoiesis (Oburoglu et al., 2014), but other lineages have not yet been examined.

Glutaminase (GLS) is an aminohydrolase that catalyzes the conversion of glutamine to glutamate, which can then be converted into the TCA cycle intermediate α- ketoglutarate (Curthoys & Watford, 1995). A potential compensatory enzyme, GLS2, is not expressed within the hematopoietic system (Heng et al., 2008). Because germline *Gls* deletion causes lethality (Masson et al., 2006), we again generated competitive bone marrow chimeras using *Gls^fl/fl^* mice (Mingote et al., 2016). Given that erythrocytes, megakaryocytes, and platelets lack CD45 expression, we generated mixed chimeras for *Gls* using β-actin green fluorescent protein (GFP) transgenic mice as wild-type competitors (**Figure 1A**). GFP expression, though minimal in mature erythrocytes, can be detected in megakaryocyte-erythroid progenitors (MEPs) and in mature platelets (Forsberg et al., 2006; Wright et al., 2001). At the end of tamoxifen treatment, the same cell subsets examined in *Cpt2*-deleted chimeras were analyzed alongside MEPs in the bone marrow and platelets in the blood (gating strategy in **Figures S1B and D**). Deletion of *Gls* was efficient (**Figure S1E**) but did not affect lymphoid or myeloid cell chimerism (**Figure 1C**). GLS-deficient MEP chimerism was slightly reduced (about 10%, p<0.05), but platelets were unaffected (**Figure 1C**). Together these data demonstrate that long-chain fatty acid oxidation and glutaminolysis are largely dispensable for lymphoid and myeloid cells, while erythroid progenitors have a partial dependence on glutaminolysis. These data suggest that other carbon sources, such as pyruvate, may be more important for hematopoiesis.

### Mitochondrial pyruvate is transiently required by both myeloid progenitors and mature cells

MPC1 and MPC2 are subunits of the mitochondrial pyruvate carrier, both of which are necessary for the transport of pyruvate into the mitochondria (Bricker et al., 2012; Herzig et al., 2012). As with *Cpt2* and *Gls*, germline deletion of either subunit of MPC is lethal (Vigueira et al., 2014). Moreover, tamoxifen-treatment of even heterozygous *Mpc2^fl/+^; ROSA26 CreER^+/-^* mice led to death within 2 weeks (data not shown), necessitating the use of mixed bone marrow chimeras. During the course of prior work on the metabolism of antibody-secreting cells (Lam et al., 2016), we preliminarily noticed that neutrophils were reduced upon *Mpc2* ablation in mixed chimeras. Other lymphoid lineages were not obviously impacted, at least at short timepoints after *Mpc2* deletion. To investigate further, we generated mixed bone marrow chimeras of wild-type CD45.1^+^ and CD45.2^+^ *Mpc2^fl/fl^; ROSA26 CreER^-/-^* or *CreER^+/-^* genotypes (McCommis et al., 2015).

A more comprehensive analysis was then performed than in our earlier results to quantify other lineages, progenitors, and the kinetics of blood lineage chimerism changes following *Mpc2* ablation.

After reconstitution, half of the chimeras were given tamoxifen and then sacrificed 10 weeks later (prolonged KO) (**Figure 2A**). This group allows for assessment of longer-term effects on lymphoid lineages, which turn over infrequently. The other half was given tamoxifen 2 weeks prior to euthanasia (recent KO). Consistent with our prior observations, the recent KO group was significantly reduced in neutrophil chimerism (**Figures 2B and C**). Ly6C^hi^ and Ly6C^lo^ monocytes were also substantially reduced immediately following *Mpc2* deletion, while DCs were slightly reduced (**Figures 2B and C**; gating strategy in **Figure S2A**). *Mpc2*-deficient neutrophils and monocytes were rescued by *Mpc2*-encoding retrovirus (**Figures S2B-D**), demonstrating that the phenotype was not due to Cre or tamoxifen toxicity. B cell chimerism was not affected in either group, whereas *Mpc2*-deficient T cell chimerism was diminished only in the prolonged KO group (**Figure 2C**). Yet contrary to our expectations, after prolonged deletion of *Mpc2*, neutrophils, Ly6C^hi^ monocytes, and Lyc6^lo^ monocytes had all recovered to mirror wild-type controls (**Figure 2C**). We considered the possibility that the myeloid recovery was driven by preferential expansion of cells that had incompletely deleted *Mpc2.* However, this seems unlikely given that this recovery would occur in competition against CD45.1^+^ wild-type cells. Indeed, *Mpc2* remained fully deleted following prolonged deletion in both the spleen and bone marrow (**Figure S2E**).

**Figure 2.**
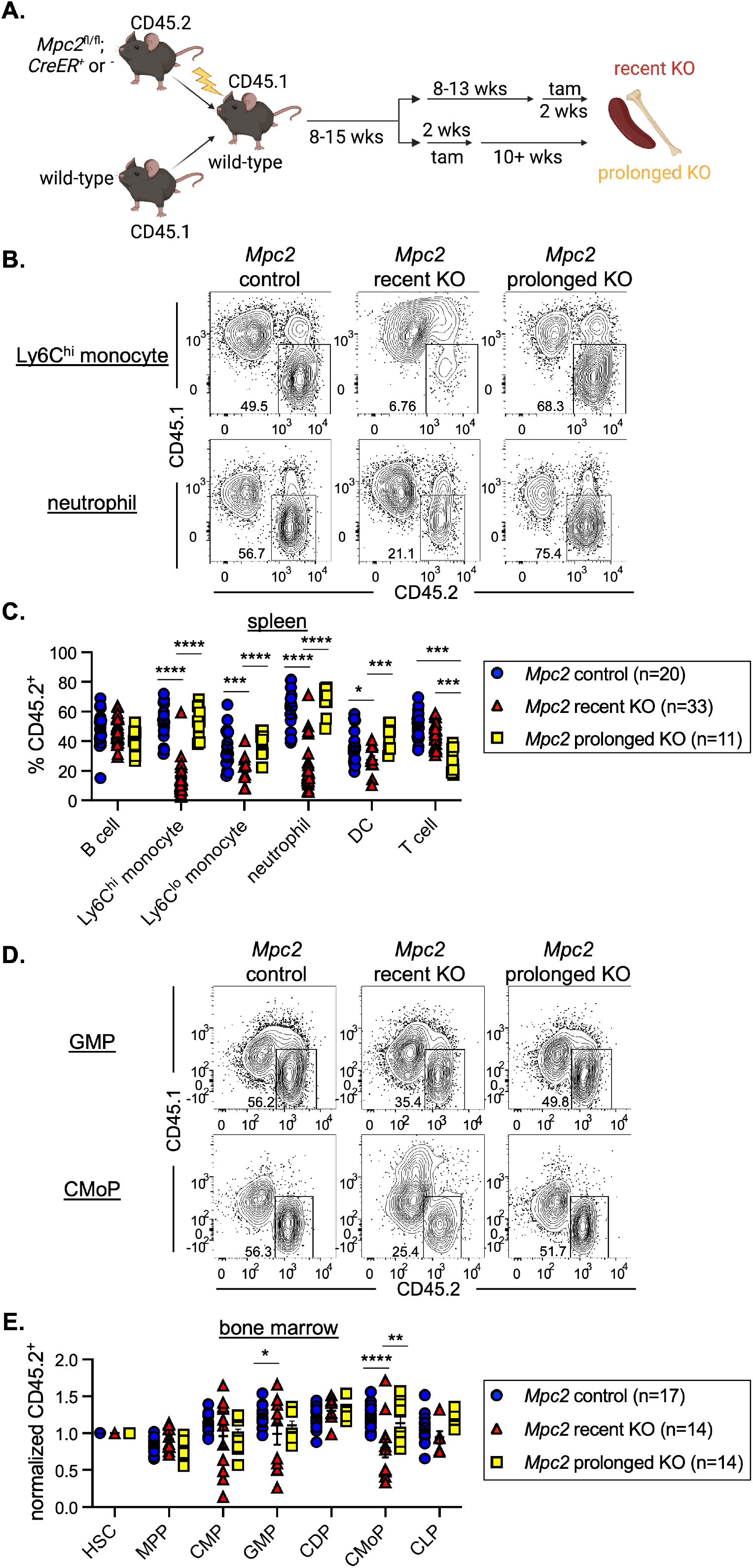
Mitochondrial pyruvate is transiently required by both myeloid progenitors and mature cells. (A) Schematic representation of mixed bone marrow chimeras to assess hematopoietic requirement for *Mpc2* (*Mpc2^fl/fl^; ROSA26 CreER^-/-^* or *ROSA26 CreER^+/-^*) both immediately following deletion and after prolonged deletion (10+ weeks post tamoxifen). (B) Representative flow cytometry plots of CD45.2 chimerism of Ly6C^hi^ monocytes and neutrophils for *Mpc2* control, recent KO, and prolonged KO chimeras. (C) CD45.2 chimerism of splenic immune lineages at early or prolonged timepoints after deletion of *Mpc2*. Each symbol represents an individual mouse. Mean values ± SEM are shown. Data are pooled from four independent experiments. *p<0.05, ***p<0.001, and ****p<0.0001 by 2-way ANOVA with post-hoc Tukey’s multiple comparisons test. (D) Representative flow cytometry plots of CD45.2 chimerism of GMPs and common monocyte progenitors (CMoPs). (E) CD45.2 chimerism of bone marrow progenitors described previously with the addition of CMoPs and common dendritic cell progenitors (CDPs) at early or prolonged timepoints after deletion of *Mpc2.* Data are normalized to HSC CD45.2 chimerism. Each symbol represents an individual mouse. Mean values ± SEM are shown, and data are pooled from three independent experiments. *p<0.05, **p<0.01, and ****p<0.0001 by 2-way ANOVA with post-hoc Tukey’s multiple comparisons test.

In the recent KO group, there was a modest reduction in *Mpc2*-deficient GMPs and a more substantial reduction in common monocyte progenitors (CMoPs) (**Figure 2D and E**; gating strategy in **Figure S2F**). CMP chimerism remained unaffected, as did that of common dendritic cell progenitors (CDPs) and CLPs (**Figure 2E**). Consistent with what was observed in mature myeloid cells, progenitor chimerism was unaffected following prolonged *Mpc2* deletion (**Figure 2E**). These results suggest that MPC2 is required for the homeostasis of the myeloid lineage, but that these cells can somehow adapt over time to the loss of mitochondrial pyruvate metabolism.

### Mature myeloid cells diminish then rapidly recover after *Mpc2* deletion

Given that myeloid cells are initially depleted upon *Mpc2* deletion but recover by 10 weeks, we performed experiments to gain a better understanding of the kinetics of this recovery. Following 8+ weeks of reconstitution, *Mpc2* chimeras were administered tamoxifen and then were bled every 2 to 3 weeks to monitor donor contribution over time relative to their pre-tamoxifen chimerism. Rather than a gradual recovery, we observed a sharp increase in *Mpc2*-deficient myeloid cells within 2 weeks after the initial nadir (**Figure 3**). These cells then further expanded beyond their pre-tamoxifen levels. In contrast, the reduction in *Mpc2*-deficient T cell chimerism was observed at 2 weeks post tamoxifen and remained relatively stable throughout the duration of the experiment (**Figure 3**). This is consistent with the requirement of MPC1 for αβ T cell development (Ramstead et al., 2020). Similarly, pyruvate dehydrogenase is also required for T cell development (Jun et al., 2021). B cells were not affected by the loss of *Mpc2* at any timepoint (**Figure 3**). We additionally examined *Cpt2* and *Gls* chimeras over time to determine if a defect would become apparent later after deletion. Yet no defects were observed (**Figures S3A and B**). These data suggest that myeloid cells are dependent on mitochondrial pyruvate import yet can adjust to the absence of MPC2 over time.

**Figure 3.**
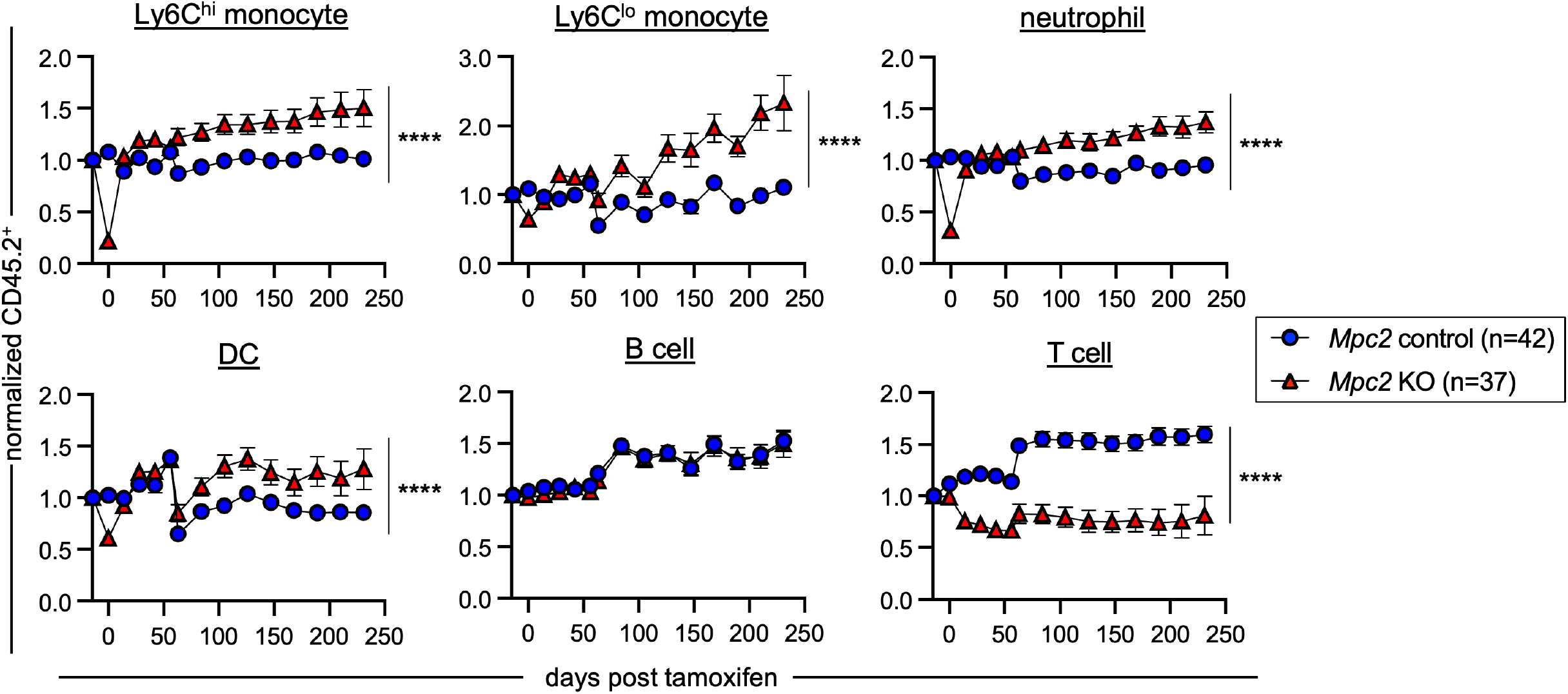
Mature myeloid cells diminish then rapidly recover after *Mpc2* deletion. CD45.2 peripheral blood chimerism of mature cell populations was assessed every 2-3 weeks in *Mpc2* chimeras. Values are normalized to pre-tamoxifen chimerism of each cell type. Data are pooled from three independent experiments. Mean values ± SEM are shown. ****p<0.0001 by paired 2-way ANOVA with post-hoc Tukey’s multiple comparisons test.

### GMPs and CMoPs proliferate rapidly immediately following *Mpc2* deletion

Given the recovery we observe in myeloid cells following the deletion of *Mpc2*, we next performed experiments to determine if MPC2-deficient cells were proliferating more and/or surviving longer to facilitate this recovery. To determine if proliferation of myeloid progenitors was altered by the loss of *Mpc2*, we injected recently deleted chimeras with bromodeoxyuridine (BrdU) 1 hour prior to sacrifice (**Figure 4A**). We then compared BrdU incorporation in the wild-type CD45.1 cells with control, recent KO, or prolonged KO CD45.2 cells within the same animals. Recent MPC2-deficient GMPs and CMoPs proliferated significantly more than did wild-type GMPs and CMoPs within the same mouse, while proliferation was similar between MPC2-deficient and -sufficient CMPs (**Figures 4B and C**). After prolonged deletion of *Mpc2*, proliferation of these populations was similar to that of their wild-type counterparts (**Figures 4B and C**). No changes were observed in the proliferation of MPC2-deficient HSCs, CLPs, or mature myeloid cells (**Figure S4A**). These data demonstrate that MPC2-deficiency triggers an adaptation in GMPs and CMoPs that is accompanied by a transient burst in proliferation. This allows for a rapid regeneration of themselves and mature myeloid populations. After prolonged *Mpc2*-deletion, this proliferative burst subsides.

**Figure 4.**
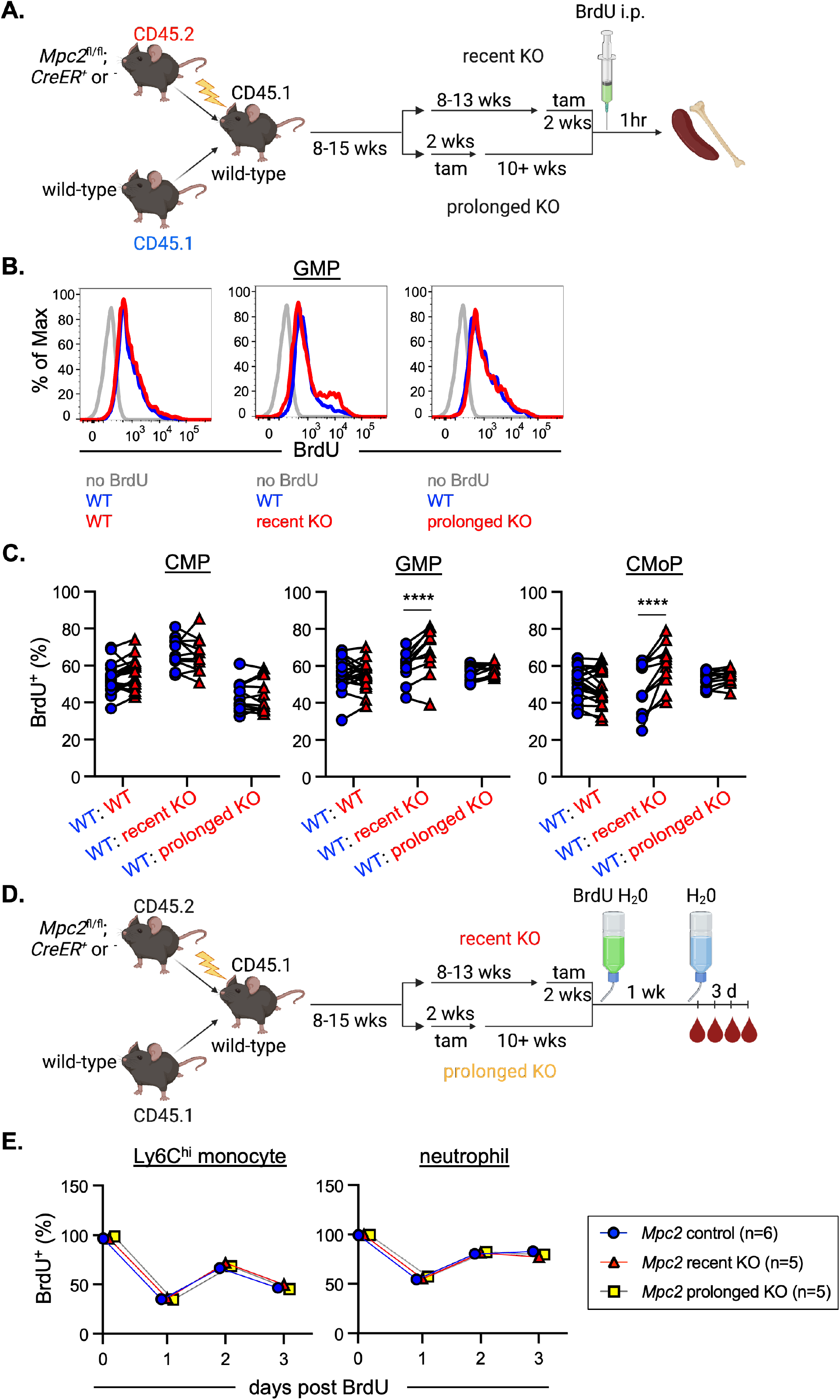
GMPs and CMoPs proliferate rapidly immediately following *Mpc2* deletion. (A) Schematic representation of BrdU pulse experiment using *Mpc2* mixed bone marrow chimeras. Chimeric mice were injected with BrdU 1 hour prior to sacrifice. (B) Representative histograms of BrdU staining in GMPs of *Mpc2* control, recent KO, and prolonged KO chimeras. Gray lines represent staining of GMPs from a mouse that was not injected with BrdU. Blue and red lines represent CD45.1^+^ (wild-type) and CD45.2^+^ (wild-type, recent KO, or prolonged KO) GMPs respectively from the same mouse injected with BrdU. (C) BrdU incorporation was measured in both CD45.1^+^ (wild-type) and CD45.2^+^ (wild-type, recent *Mpc2* deletion, or prolonged *Mpc2* deletion) CMPs, GMPs, and CMoPs following a 1-hour pulse of BrdU. Each line connects CD45.1 and CD45.2 cells within the same mouse. Data are pooled from two independent experiments. ****p<0.0001 by paired 2-way ANOVA with post-hoc Sidak’s multiple comparisons test. (D) Schematic representation of BrdU pulse-chase experiment using *Mpc2* mixed bone marrow chimeras to assess survival and turnover. Chimeras were administered BrdU water for 1 week, then were switched back to normal drinking water, at which point BrdU incorporation was assessed in peripheral blood at the time of removal from BrdU and 3 successive days afterwards. (E) BrdU incorporation was measured in peripheral CD45.2^+^ Ly6C^hi^ monocytes and neutrophils following a 3-day chase after a week of BrdU water administration. Mean values ± SEM are shown for 5-6 mice per group.

Prior studies have shown that upstream hematopoietic progenitors can sense and respond to the loss of downstream mature lineages by increasing proliferation (Boettcher & Manz, 2017; Cheshier et al., 2007; Takizawa et al., 2012). We considered the possibility that MPC2-deficient progenitors are better at sensing and proliferating in response to mature cell loss than are their wild-type counterparts. Alternatively, the initial proliferative burst in MPC2-deficient progenitors may occur entirely cell-intrinsically, perhaps due to a metabolic switch. To distinguish between these alternatives, we generated mixed bone marrow chimeras at a 90:10 ratio of wild-type to *Mpc2* KO cells. In this setting, the absolute loss in mature myeloid cells is minimal and would not be expected to trigger feedback to upstream progenitors. We observed a similar reduction in mature myeloid cells immediately following *Mpc2* deletion with a full recovery two weeks later (**Figure S4B**). This suggests that the recovery is not due to a heightened sensing of mature cell loss, but rather proliferation triggered by a cell-intrinsic metabolic switch.

To determine if the recovery of MPC2-deficient cells is also fueled by enhanced survival, we performed a BrdU pulse-chase experiment. Chimeras were administered BrdU in their drinking water for 1 week (**Figure 4D**). At the end of the week, mice were switched back to normal drinking water and were bled to establish baseline BrdU incorporation. The mice were then bled for 3 consecutive days following the cessation of BrdU administration to determine cell survival time and turnover. There were no differences observed in BrdU retention over time in peripheral Ly6C^hi^ monocytes or neutrophils between control mice or MPC2-deficient mice either immediately following deletion or after prolonged deletion (**Figure 4E**). As anticipated, B and T cell survival was also unaffected by *Mpc2* deletion (**Figure S4C**). Taken together, these data suggest that the MPC2-deficient myeloid recovery is due specifically to increased myeloid progenitor proliferation fueled by an intrinsic metabolic switch and is not due to enhanced mature cell survival.

### GMPs use glutamine *in vitro* to generate TCA cycle intermediates

The rapid proliferation of GMPs and CMoPs immediately following the loss of the MPC suggests that these progenitors switch to another carbon source to survive and expand to meet the sudden demand to produce more mature myeloid cells. To begin to define the alternate carbon source and to determine whether the switch is driven by a transcriptional adaptation, we performed RNA-sequencing (RNA-seq) on GMPs from control or *Mpc2* prolonged deletion chimeras. Only 17 genes, including *Mpc2*, were significantly dysregulated in KO GMPs (**Figure 5A**). Of note, *Bcat1* was very modestly upregulated in MPC2-deficient GMPs relative to control GMPs (about 2-fold, **Figure 5A**). BCAT1, or branched-chain amino acid transaminase 1, is a cytosolic enzyme that transfers α-amino groups from branched-chain amino acids (BCAAs; leucine, isoleucine, and valine) to α-ketoglutarate to generate α-ketoacids and glutamate (Hutson et al., 1988; Ichihara & Koyama, 1966). Overexpression of BCAT1 has been observed in both chronic and acute myeloid leukemia (Hattori et al., 2017; Raffel et al., 2017).

**Figure 5.**
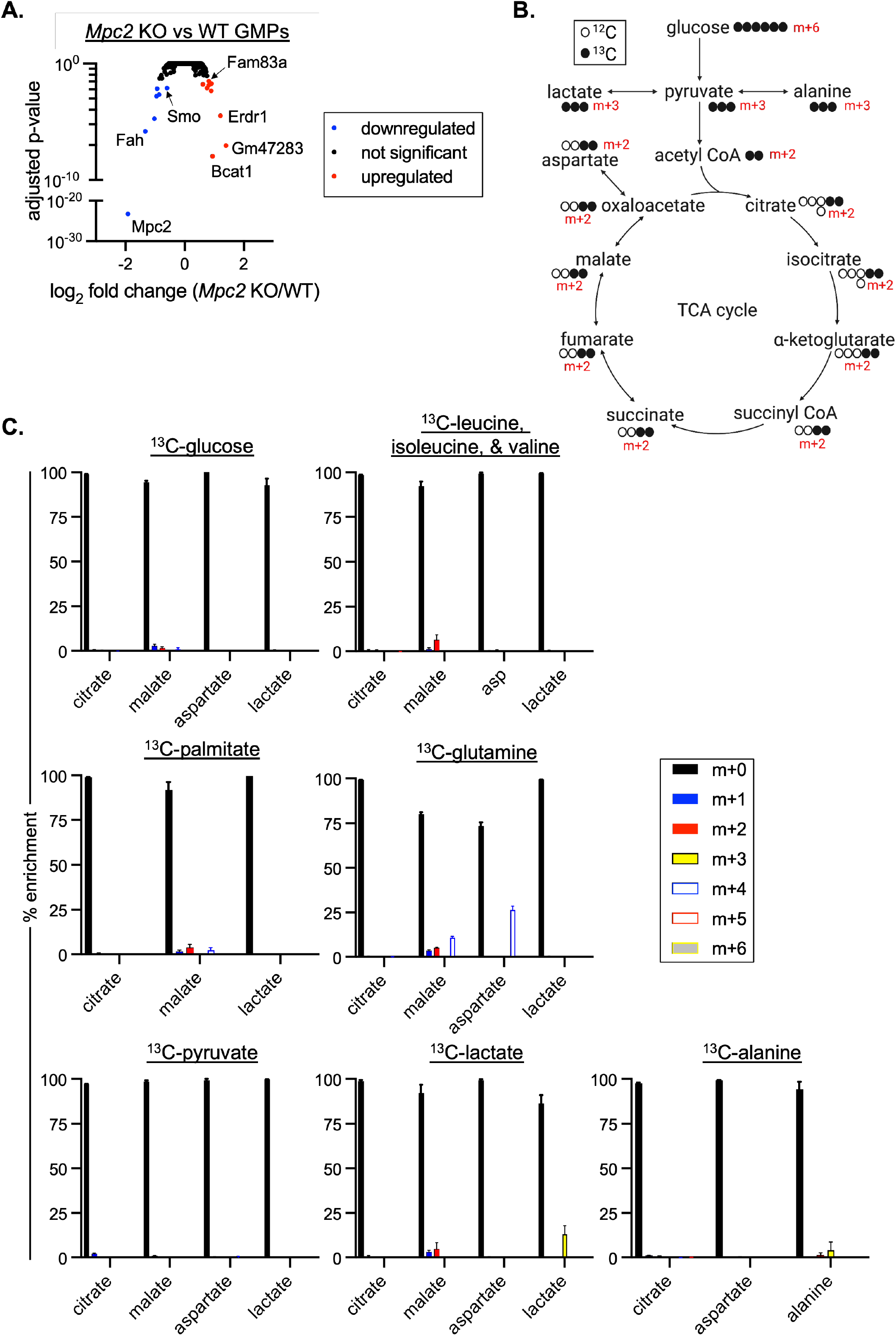
GMPs use glutamine *in vitro* to generate TCA cycle intermediates. (A) Volcano plot of gene expression fold changes between *Mpc2* prolonged KO GMPs and WT GMPs. Following RNA-seq, adjusted p-values were calculated using DESeq2. Each dot represents a gene. Five mice were analyzed for each genotype. (B) Expected outcomes of ^13^C-glucose tracing following glycolysis and mitochondrial pyruvate import. (C) LC/MS analysis of ^13^C incorporation into TCA cycle intermediates following a 24-hour culture of wild-type GMPs with uniformly ^13^C-labeled glucose, branched-chain amino acids (leucine, isoleucine, and valine), palmitate, glutamine, pyruvate, lactate, or alanine. Labeling data were corrected for natural-abundance ^13^C. Mean values ± SEM are shown for 3-5 replicates.

To begin to determine whether MPC2-deficient myeloid progenitors switch to oxidation of branched chain amino acids, we first adopted an *in vitro* differentiation assay. GMPs were sorted from wild-type mice and cultured with stem cell factor (SCF) and granulocyte-macrophage colony-stimulating factor (GM-CSF) to induce proliferation and granulocyte and myeloid differentiation. In contrast to our *in vivo* data, which show a clear requirement for mitochondrial pyruvate, treatment of cultures with the MPC inhibitor UK5099 (Halestrap & Denton, 1975), had no impact on cell survival over a 7-day period (**Figure S5A**). Similar data were observed when macrophage colony-stimulating factor (M-CSF) or granulocyte colony-stimulating factor (G-CSF) were used instead of GM-CSF (data not shown). Treatment with UK5099 did increase the amount of pyruvate excreted in the cell culture media (**Figure S5B**), demonstrating successful inhibition of MPC. These results are consistent with work showing marked differences between *in vivo* and *in vitro* metabolism in other systems (Cheng et al., 2011; Davidson et al., 2016; DeBerardinis et al., 2007; Faubert & DeBerardinis, 2017; Sellers et al., 2015). Though these *in vitro* results did not align with our *in vivo* data, the system provides an opportunity to define potential energy sources used by GMPs when mitochondrial pyruvate is not used.

To define the possible carbon sources GMPs could be using in place of pyruvate, we performed stable isotope ^13^C tracing experiments. Wild-type GMPs were cultured in media containing different ^13^C carbon sources for 24 hours, and LC/MS was used to analyze mass shifts in TCA cycle intermediates. ^13^C-glucose, which yields pyruvate through glycolysis, did not detectably contribute to the TCA cycle of *in vitro* GMPs (**Figures 5B and C**), consistent with our observations that UK5099 does not affect these cultures. We then traced ^13^C-labeled leucine, isoleucine, and valine to determine if GMPs utilize BCAAs in the TCA cycle. Labeling of TCA cycle intermediates was not observed (**Figure 5C**). In a mouse model of colon cancer, *Mpc1* ablation in intestinal stem cells leads to the preferential use of the long-chain fatty acid palmitate in intestinal crypts, while adenomas utilize glutamine (Bensard et al., 2020). To determine if GMPs could be utilizing either of these carbon sources, we followed ^13^C-labeled palmitate and glutamine. While palmitate did not detectably contribute to the TCA cycle, carbons from glutamine did contribute to the intermediates malate and aspartate (**Figure 5C**). No contributions were observed from extracellular pyruvate directly or from lactate or alanine, which are also sources of pyruvate (**Figure 5C**). Additionally, no labeling of TCA cycle intermediates was observed following culture with ^13^C-acetate or the remaining amino acids (**Figure S5C**). These data indicate that myeloid progenitors use glutamine to drive the TCA cycle when not using pyruvate, at least *in vitro*.

### Mature myeloid cells fail to fully recover after deletion of both *Mpc2* and *Gls*

To determine if myeloid progenitors switch to glutaminolysis *in vivo* when mitochondrial pyruvate is unavailable, we crossed *Gls^fl/fl^* mice to *Mpc2^fl/fl^* mice (double knockout, DKO) and established mixed bone marrow chimeras. Myeloid cells from most DKO mutant chimeras experienced the initial sharp decline we observed in MPC2- deficient chimeras (**Figure 6A**). In these chimeras, myeloid cells recovered partially (**Figure 6A**), but not to the extent observed in myeloid cells that lacked only *Mpc2* (**Figure 3**). Moreover, these results differ from those observed in cells lacking only *Gls*, which showed no competitive disadvantage to wild-type cells (**Figure 1C and S3B**). Thus, the impaired recovery in DKO cells suggests that GMPs and CMoPs switch to glutaminolysis to recover from the loss of mitochondrial pyruvate. B and T cells were also progressively reduced upon deletion of both *Gls* and *Mpc2* (**Figure 6A**). DKO GMPs and CMoPs (**Figure 6B**) were both more reduced than *Mpc2* single knockouts (**Figure 2E**). CLPs were also reduced when both *Mpc2* and *Gls* were deleted (**Figure 6B**).

**Figure 6.**
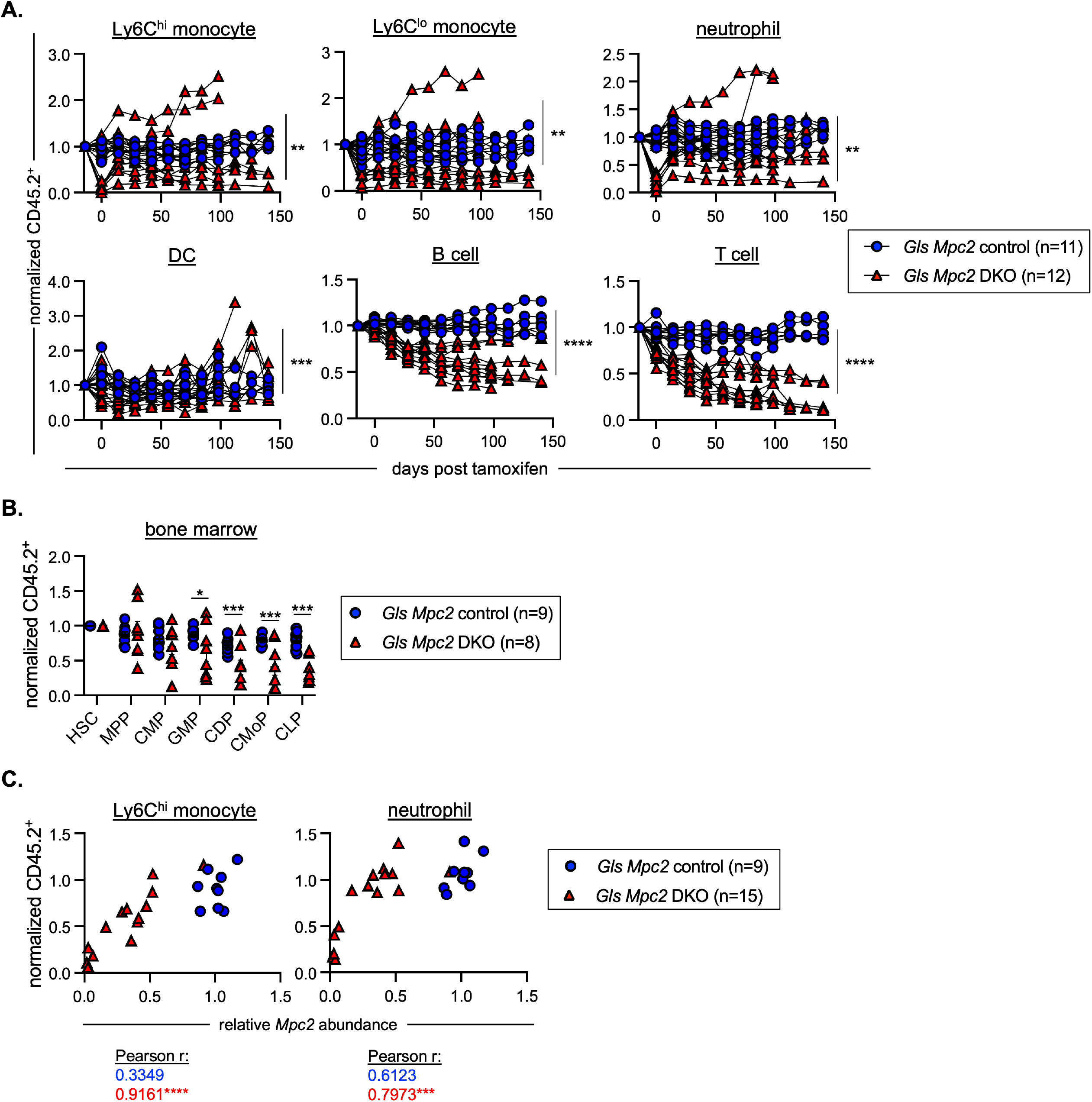
Mature myeloid cells fail to fully recover after deletion of both *Mpc2* and *Gls*. (A) Peripheral blood CD45.2 chimerism of mature cells was assessed every 2 weeks. Values are normalized to pre-tamoxifen chimerism of each cell type. Each line represents longitudinal analysis of individual mice with a symbol at each time point measured. **p<0.01, ***p<0.001, and ****p<0.0001 by paired 2-way ANOVA with post-hoc Tukey’s multiple comparisons test. (B) CD45.2 chimerism of bone marrow progenitors normalized to HSC chimerism immediately following 2 weeks of tamoxifen administration. Each symbol represents an individual mouse. Mean values ± SEM are shown. *p<0.05, ***p<0.001, and ****p<0.0001 by 2-way ANOVA with post-hoc Tukey’s multiple comparisons test. (C) Correlation between *Mpc2* expression and Ly6C^hi^ monocyte or neutrophil chimerism. Myeloid chimerism from either the blood or spleen of recent DKO mice is plotted against the *Mpc2* expression data from qPCR analysis of the CD45.2^+^ blood or spleen cells from the same mouse. Data are pooled from two independent experiments. *p<0.05, ***p<0.001, and ****p<0.0001 by Pearson correlation with 2-tailed P values.

We observed several DKO chimeras that did not display myeloid cell loss, prompting us to investigate further. Deletion of *Mpc2* was inconsistent in DKO cells, while *Gls* deletion was consistently efficient, as observed in genomic DNA from bone marrow cells and splenocytes after recent deletion (**Figure S6A**) and in RNA-seq of GMPs after prolonged deletion (**Figure S6B**). The extent of remaining *Mpc2* correlated well with

Ly6C^hi^ monocyte or neutrophil chimerism (**Figure 6C**). Given the efficiency of deletion of *Mpc2* in single mutant cells (**Figures S2E and S6B**), it seems unlikely that the floxed allele is inherently difficult for Cre to access or recombine. Rather, these data suggest a selection against cells that have fully deleted both *Mpc2* and *Gls*.

In the setting of competitive mixed chimeras, such selection events would be predicted to result in a pronounced reduction in chimerism unless they were to take place gradually and allow time for incompletely deleted cells to compensate. Similarly, a gradual deletion of *Mpc2* may allow time for cells to switch to glutaminolysis and recover, whereas a rapid loss of *Mpc2* would not allow for such a switch and myeloid recovery. To test these possibilities, we compared the kinetics of *Mpc2* deletion in *CreER^+/+^* and *CreER^+/-^* chimeras. *CreER^+/+^* cells showed substantial deletion of *Mpc2* in most mice between days 3 and 6 of tamoxifen administration, whereas in *Cre^+/-^* cells, this was not observed until days 9 through 12 (**Figure S6C**). This more rapid loss of *Mpc2* in *Cre^+/+^* cells was accompanied by a significantly slower myeloid recovery (∼30 weeks) relative to *CreER^+/-^* cells (∼2 weeks; **Figure S6D**). These results are consistent with a mechanism by which subtle or gradual metabolic perturbations, as would be expected under normal dietary and circadian conditions, allows for progenitors to tune catabolic programs to adapt and maintain homeostasis.

## Discussion

Hematopoietic stem and progenitor cells maintain homeostasis by matching blood lineage replenishment to the rate of mature cell loss. A failure to maintain homeostasis can lead to myeloproliferative disorders and cancer, lymphopenia and susceptibility to infection, and anemia from reduced red blood cells. To maintain homeostasis, stem and progenitor cells must sense and respond to fluctuations in mature cell numbers, ranging from a significant loss of cells due to acute insults such as infections and to minor daily changes such as metabolic perturbations. The extreme loss of mature cells triggers the proliferation and differentiation of progenitors (Bhattacharya et al., 2006; Cheshier et al., 2007), which is practical for acute insults. However, during steady state, an anticipatory mechanism would be preferable to adjust progenitor output and prevent the dramatic loss of mature cells.

Under steady-state conditions, a switch from glycolysis to oxidative phosphorylation promotes ATP production to support transit amplification (Takubo et al., 2013; Yu et al., 2012; Wang et al., 2014; Maryanovich et al., 2015; Ito & Suda, 2014; Nakamura-Ishizu et al., 2020; Simsek et al., 2010; Suda et al., 2011; Ho et al., 2017). Altering this metabolic switch can have severe consequences for HSCs and/or differentiated cells (Inoue et al., 2010; Ito et al., 2004; Norddahl et al., 2011; Tothova et al., 2007; Wang et al., 2014; Yu et al., 2012). The specific carbon sources that fuel oxidative phosphorylation in hematopoietic progenitors *in vivo* have not been defined.

Here, we demonstrated that the genetic ablation of pyruvate import into the mitochondria sharply reduced mature myeloid cells, but these cells recovered rapidly due to a transient proliferative burst specifically by myeloid progenitors. This recovery of MPC2-deficient myeloid cells occurred in competition with wild-type cells, demonstrating that this proliferative burst was intrinsic to cells devoid of mitochondrial pyruvate import. The progenitor proliferation was only transient and did not result in MPC2-deficient myeloid cells substantially outcompeting their wild-type competitors. These findings are consistent with prior studies showing relatively stable numbers of circulating mature cells despite marked changes in nutrient availability, altered by circadian rhythms and diet (Pickel & Sung, 2020; Potter et al., 2016; Schlierf & Dorow, 1973). These data further highlight the precise control progenitors have in maintaining homeostasis.

This proliferative response and metabolic adaptation to the absence of mitochondrial pyruvate is not unique to myeloid progenitors. Schell et al. demonstrated that in Drosophila, deletion of *Mpc1* increases intestinal stem cell proliferation (Schell et al., 2014). In a mouse model of colon cancer, the deletion of *Mpc1* in intestinal stem cells induces a preferential switch to glutamine in *ex vivo* adenomas and leads to increased tumor burden *in vivo* (Bensard et al., 2020). Additionally, MPC1-deficient mouse embryonic fibroblasts utilize increased glutaminolysis relative to wild-type controls (Vanderperre et al., 2016).This switch to glutamine following inhibition of MPC has also been demonstrated in various cancer cell lines (Yang et al., 2014). While our data likewise indicated this proliferative response induced by a metabolic switch, we found that this burst was only transient, leading to rapid restoration of homeostasis but no myeloproliferative disorders. One possible mechanism is that MPC2-deficient progenitors briefly switch to lactate production due to increased cytosolic pyruvate. This has been shown to occur in tumors as part of the Warburg effect, leading to regeneration of NAD^+^, which in turn enables repeated cycles of glycolysis and biomass production to enable rapid proliferation (DeBerardinis et al., 2008; Vander Heiden et al., 2009; Warburg, 1925). Yet once MPC2-deficient progenitors switch to glutaminolysis, feedback mechanisms such as citrate-mediated inhibition of phosphofructokinase (PFK) activity (Garland et al., 1963; Poorman et al., 1984) might shut down this increased glycolysis and normalize the proliferative rate. These changes are difficult to directly observe *in vitro*, as demonstrated by our experiments in which mitochondrial pyruvate was minimally utilized even by untreated wild-type myeloid progenitors. Yet consistent with this possible mechanism, deletion of lactate dehydrogenase A (LDHA) permanently impairs proliferation of hematopoietic progenitors *in vivo*, as does loss of the M2 isoform of pyruvate kinase (PKM2), which limits the initial formation of pyruvate (Wang et al., 2014).

When we ablated both *Gls* and *Mpc2*, the recovery of mature myeloid cells was significantly impaired, despite the fact that deletion of *Gls* alone had no discernable effect on myeloid cells. This suggests that glutaminolysis is functionally required only when mitochondrial pyruvate is unavailable. Yet there might be additional carbon sources that promote recovery as well. For example, while MPC1-deficiency in mice leads to lethality around E13.5, administering pregnant dams a ketogenic diet restores embryonic development (Vanderperre et al., 2016). Additionally, ablation of *Mpc1* in the intestinal epithelium of mice led to increased palmitate oxidation relative to wild-type controls in *ex vivo* intestinal crypts (Bensard et al., 2020). Similarly, although myeloid progenitors primarily switch to glutaminolysis in the absence of mitochondrial pyruvate, we have previously observed that MPC2-deficient neutrophils can be rescued by retroviral transduction with a long-chain fatty acid transporter (Lam et al., 2016; Schaffer & Lodish, 1994). Although we did not observe contributions from palmitate *in vitro*, preliminary experiments showed a selection against cells that had deleted both *Cpt2* and *Mpc2* in double knockout chimeras (data not shown). These data suggest a possible role for long-chain fatty acid oxidation to complement MPC2-deficiency *in vivo*. Additionally, while we did not trace carbons from BCAAs to TCA cycle intermediates *in vitro*, RNA-seq revealed an upregulation of *Bcat1* in MPC2-deficient GMPs relative to wild-type, suggesting BCAAs may also utilized.

Regardless of the metabolic switch that is occurring in the absence of mitochondrial pyruvate, progenitors clearly need time to adjust then proliferate and differentiate enough to recover mature myeloid cells. By altering the rate of *Mpc2* ablation, we demonstrated that mature myeloid cells did not recover as quickly when deletion occurred rapidly as they did after more gradual deletion. This observation suggests that myeloid progenitors can sense and respond to the gradual loss of mitochondrial pyruvate. A rapid loss of mitochondrial pyruvate, however, results in insufficient time for progenitors to enact a metabolic switch, survive, and mediate recovery. This slower deletion would seem to be a more physiologically relevant model of nutrient fluctuations, as subtle or gradual metabolic perturbations would be expected under normal dietary and circadian conditions as opposed to the sudden loss of a carbon source.

Taken together, these data demonstrate that hematopoietic progenitors can intrinsically respond in real time to the anticipated loss of their daughter cells. In the example we provide here, this adaptation is triggered by a metabolic switch that more dramatically impacts mature cells than the upstream progenitors. This ability to sense and respond to perturbations allows progenitors to adapt to changes in progeny turnover rates without necessarily waiting for feedback caused by extreme mature cell loss. These data help explain how the hematopoietic system buffers against marked fluctuations in blood cells to maintain homeostasis.

## Acknowledgements

This work was funded by the National Institutes of Health grants R01AI129945 (D.B.), R35ES2028365 (G.J.P.), and R01DK116746 (M.J.W.). Development of *Mpc2* floxed mice was supported by NIH R01DK104735 (B.N.F.).

## Author Contributions

Conceptualization, H.A.P. and D.B.; Methodology, H.A.P., Y.W., G.J.P., and D.B.; Formal Analysis, H.A.P., Y.W. G.J.P., and D.B.; Investigation, H.A.P. and Y.W.; Resources, M.J.W., B.N.F., G.J.P., and D.B.; Writing – Original Draft, H.A.P. and D.B.; Writing – Reviewing & Editing, H.A.P., Y.W., M.J.W., B.N.F., G.J.P., and D.B.; Supervision, G.J.P. and D.B.; Project Administration, D.B.; Funding Acquisition, M.J.W., B.N.F., G.J.P., and D.B.

## Methods

### Mice

All animal procedures used in this study were approved by the Animal Care and Use Committees at Washington University in St. Louis and the University of Arizona. C57BL6/N and B6.Ly5.2 (CD45.1) mice were obtained from Charles River Laboratories.

β-actin GFP transgenic mice were obtained from The Jackson Laboratory (stock number 006567). *Mpc2^fl/fl^* mice were kindly provided by Dr. Brian N. Finck (Washington University in St. Louis) and were generated as described (McCommis et al., 2015) (currently available from The Jackson Laboratory, stock number 032118). *Cpt2^fl/fl^* mice were kindly provided by Dr. Michael J. Wolfgang (Johns Hopkins) and were generated as described (Lee et al., 2015). *Gls^fl/fl^* mice were generated as described (Mingote et al., 2016) and were obtained from The Jackson Laboratory (stock number 017894). All floxed mouse lines were crossed to *ROSA26 CreER* mice obtained from The Jackson Laboratory (stock number 008463).

### Mixed Bone Marrow Chimeras

To generate mixed bone marrow chimeras, bone marrow cells from a CD45.2^+^ donor mouse carrying floxed alleles were mixed 1:1 or 1:9 with bone marrow cells from B6.Ly 5.2 CD45.1 mice (or β-actin GFP transgenic mice for *Gls* chimeras). 10 x 10^6^ total mixed bone marrow cells were injected retro-orbitally into 800 cGy-irradiated B6.Ly 5.2 CD45.1 mice. Recipient mice were administered Sulfatrim (40mg/mL sulfamethoxazole and 8mg/mL trimethoprim) via drinking water for 2 weeks. Chimeras were rested for at least 8 weeks to allow reconstitution. Peripheral blood chimerism was assessed through tail venipuncture prior to tamoxifen administration. To induce deletion, tamoxifen was administered in their diet (400mg tamoxifen citrate/kg diet, Envigo) for 2 weeks. *Mpc2* retroviral chimeras were generated as previously described (Lam et al., 2016).

### Mouse Tissue Processing

Spleens were harvested and dissociated using frosted glass microscope slides. Femurs, tibiae, and pelvic bones were isolated and crushed with a mortar and pestle. Red blood cells were lysed using a 0.15M NH4Cl, 10mM KHCO3, 0.1mM EDTA, pH 7.2 solution (ACK). Non-cellular debris was removed by gradient centrifugation for 10 minutes at 2000g using Histopaque 1119 (Sigma-Aldrich). Interface cells were collected and washed, then filtered through 70μm nylon mesh prior to antibody staining or downstream applications. Peripheral blood was collected in 10mM EDTA/phosphate buffered saline (PBS) via tail venipuncture of warmed mice. Red blood cells were lysed with ACK prior to antibody staining. For platelets, immediately following bleeding and prior to ACK lysis, 2% dextran sulfate/PBS was added to blood samples. Samples were centrifuged at 200g for 20 minutes at room temperature. The supernatant was transferred to a new tube and centrifuged at 1100g for 10min at room temperature to pellet platelets.

### Flow Cytometry

Single cell suspensions were prepared from spleen, bone marrow, or peripheral blood. Cells were resuspended in PBS with 5% adult bovine serum and 2mM EDTA prior to staining. The following antibodies were purchased from BioLegend: CD3 (clone 17A2) – PE/Dazzle 594, Brilliant Violet 510, PerCP/Cy5.5; B220 (RA3-6B2) – Alexa Fluor 700, APC, Brilliant Violet 421, FITC; I-A/I-E (M5/114.15.2) – Biotin, Brilliant Violet 711; Ly-6C (HK1.4) – Brilliant Violet 510, FITC; Ly-6G (1A8) – Brilliant Violet 785, FITC, PE; CD45.1 (A20) – APC/Cyanine 7, Brilliant Violet 605, PE/Cy7; CD45.2 (104) – APC/Cy7, Brilliant Violet 421, Brilliant Violet 510; NK-1.1 (PK136) – PE; IgD (11-26c.2a) – Brilliant Violet 605; CD90.1 (OX-7) – FITC; CD115 (AFS98) – Brilliant Violet 711; CD127 (A7R34) – Biotin; CD138 (281-2) – Brilliant Violet 650, Brilliant Violet 510; CD150 (TC15-12F12.2) – Brilliant Violet 421, Alexa Fluor 647; IgM (RMM-1) – APC; Ly-6A/E (E13-161.7) – PE; Ly- 6G/Ly-6C (RB6-8C5) – APC; Streptavidin – APC/Cy7, Brilliant Violet 650, Brilliant Violet 711; CD61 (2C9.G2 (HMβ3-1)) – PE; CD8 (53-6.7) – Alexa Fluor 700; CD11b (M1/70) – Alexa Fluor 488, PerCP/Cy5.5; CD11c (N418) – APC/Cyanine7, Biotin; CD16/32 (93) – PerCP/Cyanine5.5; CD34 (MEC14.7) – Brilliant Violet 421. The following antibodies were purchased from eBioscience: CD115 (AFS98) – Biotin; Ly-6G/Gr-1 (RB6-8C5) – PE-Cy5; CD4 (GK1.5) – PE-Cy5; CD8 (53-6.7) – PE-Cy5; CD11b (M1/70) – PE-Cy5; CD27 (LG.7F9) – APC; CD34 (RAM34) – FITC. The following antibodies were purchased from BD: CD11b (M1/70) – BUV395, BUV661, PE-Cy7; CD4 (RM4-5) – PE-Cy7; CD8 (53-6.7) – BUV 805; CD135 (A2F10.1) – PECF594; Streptavidin – BV421, BV605; CD3 (clone 17A2) – PE; CD27 (LG.3A10) – BUV805; CD34 (RAM34) – BV786. The following antibodies were purchased from Invitrogen: CD11c (N418) – PerCP-Cyanine5.5; CD115 (AFS98) – APC; IgM (II/41) – PerCP-eFluor 710, FITC; CD117 (2B8) – PE-Cyanine7; TER-119 (TER-119) – PE-Cyanine7; CD4 (RM4-5) – Pacific Orange; CD16/32 (93) – APC; CD34 (RAM34) – Alexa Fluor 700. DAPI and propidium iodide were purchased from Sigma-Aldrich. Cells were analyzed on a BD LSR II, BD Fortessa, BD Fortessa X-20, or Cytek Aurora. All fluorescence activated cell sorting was performed on a BD FACS Aria II or III. Data was analyzed using FlowJo software (FlowJo Enterprise).

### Quantitative PCR Analysis

To quantify deletion of *Cpt2*, *Gls*, and *Mpc2*, 50,000 to 150,000 CD45.2^+^ cells were sorted from the spleen, bone marrow, or peripheral blood of chimeras into PBS with 5% adult bovine serum. DNA was isolated using the Genomic DNA Mini Kit (IBI Scientific). Quantitative PCR was then used to determine the extent of deletion. Normalization control primers amplified a region of GAPDH (forward primer: 5’-AACTTTGGCATTGTGGAAGG- 3’ and reverse primer 5’-GGATGCAGGGATGATGTTCT-3’). Primers for *Cpt2*, *Gls*, and *Mpc2* amplified a region within the loxP sites. The *Cpt2* forward primer was 5’- GATGGCTGAGTGCTCCAAAT-3’, and the reverse was 5’- GCCAGACCCAAGGTGTTCT-3’. The *Gls* forward primer was 5’- ATCTCCTTGCCCTCGCTGT-3’, and the reverse was 5’-CGCCCTCGGAGATCCTAC-3’. The *Mpc2* forward primer was 5’-CCCCACCGTGTCTTAATGTC-3’, and the reverse was 5’-CACAGTGGACTGAGCTGTGC-3’. PCR reactions were performed in 20μL containing SYBR Green PCR Master Mix (Applied Biosystems) and 10μM of each primer. Reactions were performed in a StepOnePlus realtime qPCR machine (Applied Biosystems) with 40 cycles of the following program 94°C-30 seconds, 59.1°C-30 seconds, 72°C-30 seconds, followed by a melt-curve analysis. Quantification was calculated using normalized ΔCt values. An average from wild-type samples was used to estimate a value of 1.

### BrdU Pulse and Pulse-Chase Experiments

For BrdU pulse experiments, mice were injected with 1mg BrdU intraperitoneally 1 hour prior to sacrifice. For BrdU pulse-chase experiments, mice were provided with 0.8mg/mL BrdU in their drinking water replaced daily for 7 days. At the end of the 7 days, mice were returned to normal drinking water and were bled that day and the following 3 days. Cells were then processed and stained for BrdU incorporation with the FITC BrdU Flow Kit (BD) per manufacturer’s instructions.

### RNA-sequencing

CD45.2^+^ GMPs were sorted from wild-type (*ROSA26 CreER^+/-^*), *Mpc2*, *Gls*, and *Gls Mpc2* KO chimeras about 10 weeks post tamoxifen treatment into Buffer RA1 (Macherey-Nagel). RNA was isolated using the NucleoSpin RNA XS kit (Macherey-Nagel). Sequencing libraries were generated by Novogene using the SMARTseq V4 kit (Takara Bio). Paired end 150bp reads were acquired by Novogene using Illumina NovaSeq. For quantification of differences in gene expression, reads were mapped to Gencode’s M27 annotation files, and transcript abundances were calculated using Salmon (Patro et al., 2017). DESeq2 was used to quantify differentially expressed genes (Love et al., 2014). To analyze exon usage, fastq files were first aligned to the mm10 genome using HISAT2 (Kim et al., 2015). HISAT2 bam files were then visualized using Integrative Genomics Viewer (IGV) (Thorvaldsdóttir et al., 2013) and used to quantify transcripts per kilobase million (TPM) for floxed exons via DEXSeq (Anders et al., 2012).

### GMP Cultures

Single cell suspensions of bone marrow cells were generated as per the tissue processing section. C-kit^+^ cells were enriched using CD117 microbeads (Miltenyi Biotec). GMPs were sorted into RPMI containing 10% fetal bovine serum (FBS), glutamine, and penicillin/streptomycin. 50,000 to 100,000 GMPs were cultured for pyruvate excretion analysis or ^13^C tracing experiments, while 1,000 GMPs were plated for 7-day differentiation analysis, in RPMI containing 10% FBS, glutamine, penicillin/streptomycin, 10ng/mL GM-CSF (PeproTech), and 10ng/mL SCF (PeproTech). Cells were incubated at 37°C with 5% CO2.

### Nutrient-uptake Analysis

GMPs were sorted and cultured as described above. DMSO or 10μM UK5099 (Sigma Aldrich) was added to the medium as a vehicle control or to inhibit MPC. GMPs were collected on day 3 of culture, washed with PBS, replated in fresh media, and returned to the incubator for 48 hours. The spent media was collected. The extraction solvent methanol/acetonitrile (ACN) /water (2:2:1) containing isotope-labeled internal standards ([U-^13^C6]glucose; [U-^13^C3]lactate; [U-^13^C5]glutamine; and [U-^13^C5]glutamate; Cambridge Isotope Laboratories) were spiked into media samples. Samples were vortexed for 30 seconds, incubated for 1 minute in liquid nitrogen, and then sonicated for 10 minutes. Following a 1-hour incubation at -20°C, the samples were centrifuged at 14,000rpm for 10 minutes. The supernatant was analyzed by using an Agilent 1290 UHPLC system coupled to an Agilent 6540 quadruple time-of-flight (Q-TOF) mass spectrometer. The consumption rates (*x*) were normalized by cell growth over the experimental time period using the following algorithm (*N0*, starting cell number; *t*, incubation time; *DT*, doubling time; *Y*, nutrient utilization).

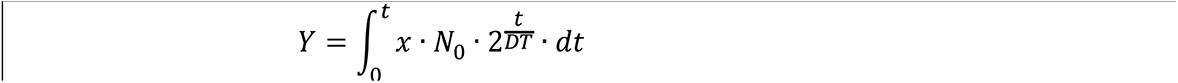

### Isotope Labeling and Metabolite Extraction

GMPs were sorted and cultured as described above. Cells were collected on day 4 of culture, washed with PBS, and replated in the appropriate media (listed below) for 24 hours. Following incubation, cells were collected, washed with PBS, washed with HPLC-grade water, and quenched with HPLC-grade methanol. Pellets were dried using a SpeedVac. Dried cell pellets were extracted with 1mL methanol/ACN/water (2:2:1). Samples were vortexed for 30 seconds, incubated for 1 minute in liquid nitrogen, and then sonicated for 10 minutes. This process was repeated three times. After a 1-hour incubation at -20°C, samples were centrifuged at 14,000rpm for 10 minutes. The supernatant was collected and dried via SpeedVac and reconstituted in ACN/water (1:1) for LC/MS analysis.

All U-^13^C carbon sources were obtained from Cambridge Isotope Laboratories. For all ^13^C experiments, the base medium was RPMI containing 10% FBS, glutamine, penicillin/streptomycin, 10ng/mL GM-CSF, and 10ng/mL SCF, unless noted otherwise, and supplemented with ^13^C carbon sources. For glucose, glucose-free RPMI was used to make the base medium instead of standard RPMI and was supplemented with 11.1mM ^13^C-glucose. For glutamine, glutamine-free RPMI was used, glutamine was left out of the base medium, and 4mM ^13^C-glutamine was added. For palmitate, 100μM ^13^C-palmitate-BSA and 100μM oleate-BSA was added. For pyruvate, 100μM ^13^C-pyruvate was supplemented. Concentrations used for all other ^13^C tracing experiments were as follows: 3mM ^13^C-lactate, 400μM ^13^C-alanine, 500μM ^13^C-acetate, 430μM ^13^C-asparagine, 382μM ^13^C-leucine, 382μM ^13^C-isoleucine, 171μM ^13^C-valine, 220μM ^13^C-lysine, 91μM ^13^C-phenylalanine, 60μM ^13^C-tryptophan, 110μM ^13^C-tyrosine, 336μM ^13^C-cysteine, 133μM ^13^C-glycine, 286μM ^13^C-serine, 136μM ^13^C-glutamate, 1.15mM ^13^C-arginine, 97μM ^13^C- histidine, 174μM ^13^C-proline, 101μM ^13^C-methionine, 168μM ^13^C-threonine, 91μM ^13^C- phenylalanine, 110μM ^13^C-tyrosine, and 150μM ^13^C-aspartate.

### LC/MS Analysis

Media samples and cell extracts were analyzed with Agilent 1290 UPLC system coupled to an Agilent 6540 quadruple time-of-flight (Q-TOF) mass spectrometer. Polar metabolites were separated on a zic-pHILIC column (100mm × 2.1mm, 5μm, polymer; Merck-Millipore) with a ZIC-pHILIC guard column (20mm × 2.1mm, 5μm, polymer; Merck-Millipore). The column compartment temperature was 40°C. Mobile phase A was 95% water and 5% ACN with 20mM ammonium bicarbonate, 0.1% ammonium hydroxide solution, and 2.5μM medronic acid. The ammonium hydroxide solution was purchased as 25% ammonia in water (Honeywell). Mobile phase B was 95% ACN and 5% water with 2.5μM medronic acid. Separation was carried out with a flow rate of 250μL minutes^-1^ and the following linear gradients: 90% B from 0 to 1 minutes, 90% to 25% B from 1 to 14 minutes, 25% B from 14 to 15.5 minutes, and 25% B to 90% B from 15.5 to 18 minutes. The column was equilibrated with 90% B for 10.5 minutes at a flow rate of 400μL minutes^-1^ and 1.5 minutes at a flow rate of 250μL minutes^-1^. Mass spectrometry detection was accomplished in negative ionization mode with the following settings: gas temperature 200°C, sheath gas temperature 300°C, drying gas flow rate 10L minutes^-1^, sheath gas flow rate 12L minutes^-1^, nebulizer pressure 44psi, capillary voltage 3000V, nozzle voltage 2000V, fragmentor voltage 100V, skimmer voltage 65V, and scan rate 1 spectrum second^-1^.

### Statistics

All statistical analyses were conducted using GraphPad Prism and are detailed in each figure legend.

## Figures

Figures were created with BioRender.com.

## Supplemental Figure Titles and Legends

**Figure S1.**
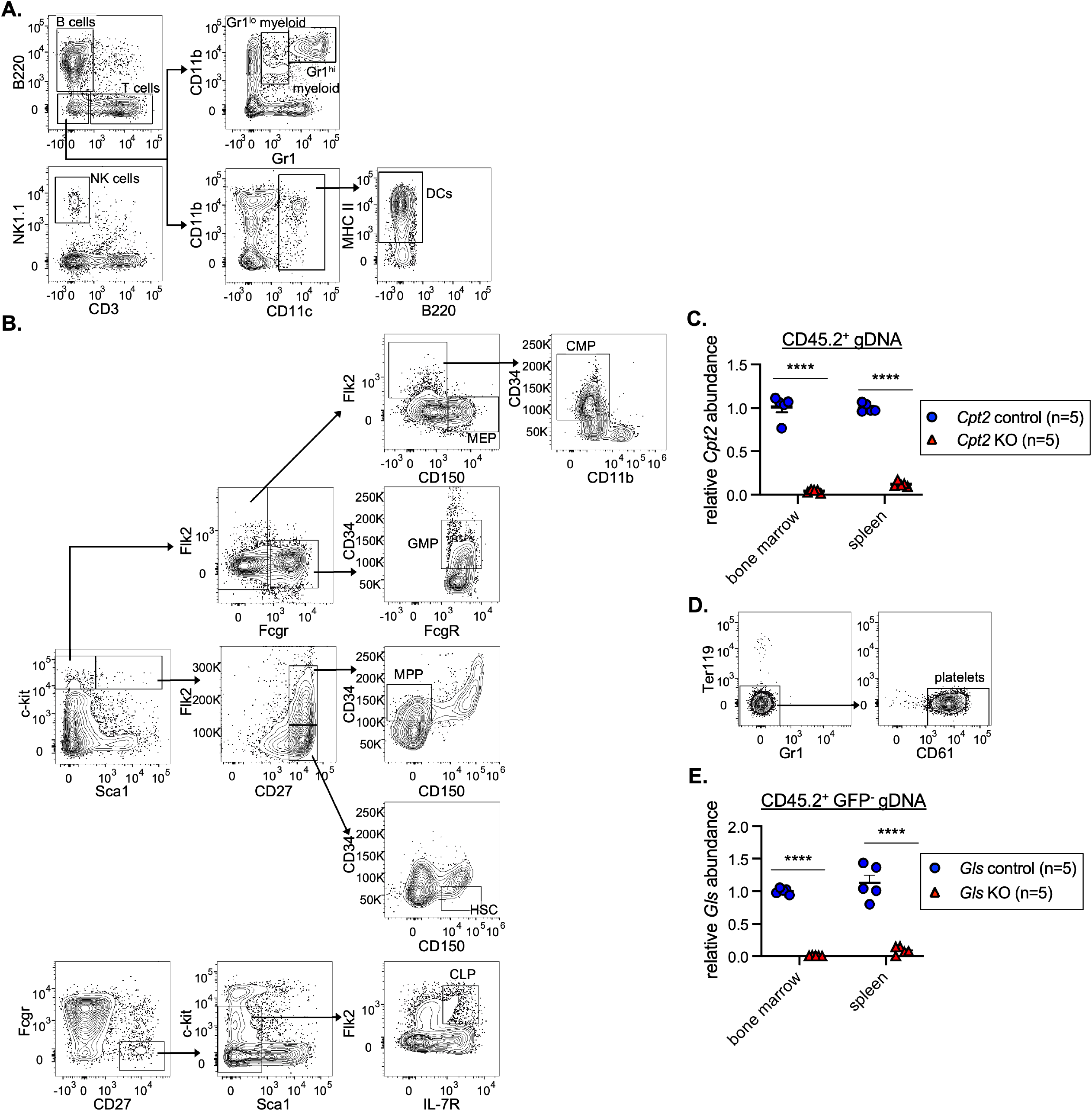
related to Figure 1. Gating strategy and quantification of deletion of *Cpt2* and *Gls*. (A) Representative flow cytometry gating strategy for splenic populations. (B) Representative flow cytometry gating strategy for bone marrow populations. (C) Quantitative PCR analysis of genomic *Cpt2* deletion. CD45.2^+^ cells from the bone marrow or spleen were sorted for genomic DNA extraction. DNA quantification within loxP sites was quantified relative to wild-type cells and GAPDH. Mean values ± SEM are shown. ****p<0.0001 by Student’s 2-tailed t-test. (D) Representative flow cytometry gating strategy for platelets in the blood. (E) Quantitative PCR analysis of genomic *Gls* deletion. CD45.2^+^ GFP^-^ cells from the bone marrow or spleen were sorted for genomic DNA extraction. DNA quantification within loxP sites was quantified relative to wild-type cells and GAPDH. Mean values ± SEM are shown. ****p<0.0001 by Student’s 2-tailed t-test.

**Figure S2.**
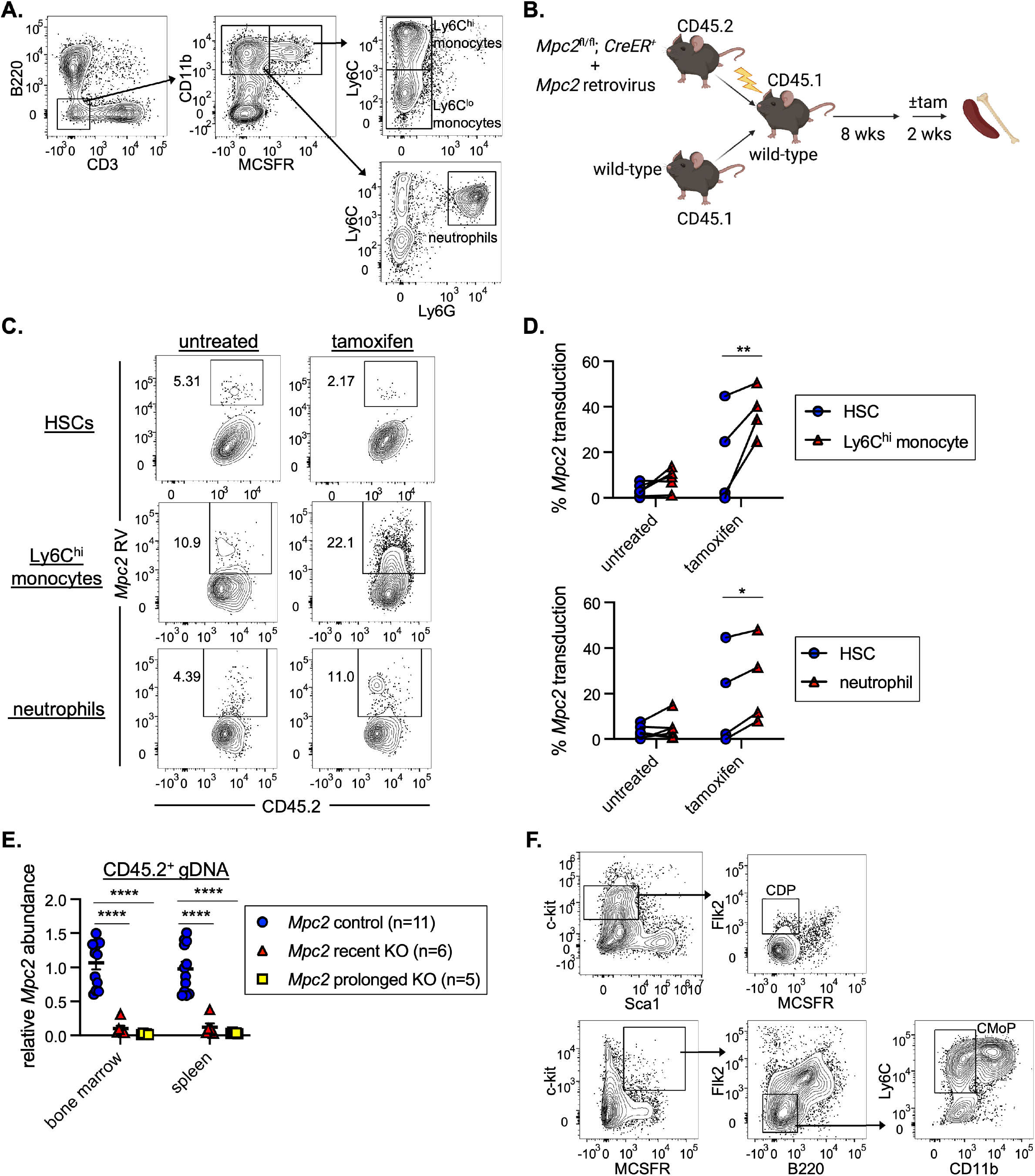
, related to Figure 2. Gating strategy, quantification of deletion of *Mpc2*, and *Mpc2* retroviral rescue of MPC2-deficient myeloid cells. (A) Representative flow cytometry gating strategies for splenic Ly6C^hi^ and ^lo^ monocytes and neutrophils. (B) Schematic representation of *Mpc2* retroviral bone marrow chimera experiments to assess ability to rescue *Mpc2* KO (+ tam) or control (no tam) Ly6C^hi^ monocytes and neutrophils. C-kit^+^ cells were enriched from the bone marrow of *Mpc2^fl/fl^; ROSA26 CreER^+/-^* mice and transduced with *Mpc2* retrovirus prior to transplantation alongside bone marrow from wild-type mice into irradiated recipients. (C) Representative flow cytometry plots of *Mpc2* retrovirus (RV) expression in HSCs, Ly6C^hi^ monocytes, and neutrophils of an individual control or KO mouse. (D) *Mpc2* transduction in HSCs paired with Ly6C^hi^ monocytes (top) or neutrophils (bottom) from the same mouse. *p<0.05 and **p<0.01 by Student’s 2-tailed paired t-test. (E) Quantitative PCR analysis of genomic *Mpc2* deletion in *Mpc2* chimeras. CD45.2^+^ cells from the bone marrow or spleen were sorted for genomic DNA extraction. DNA quantification within loxP sites was quantified relative to wild-type cells and GAPDH. Mean values ± SEM are shown. ****p<0.0001 by 1-way ANOVA with post-hoc Tukey’s multiple comparisons test. (F) Representative flow cytometry gating strategies for CDPs and CMoPs in the bone marrow.

**Figure S3.**
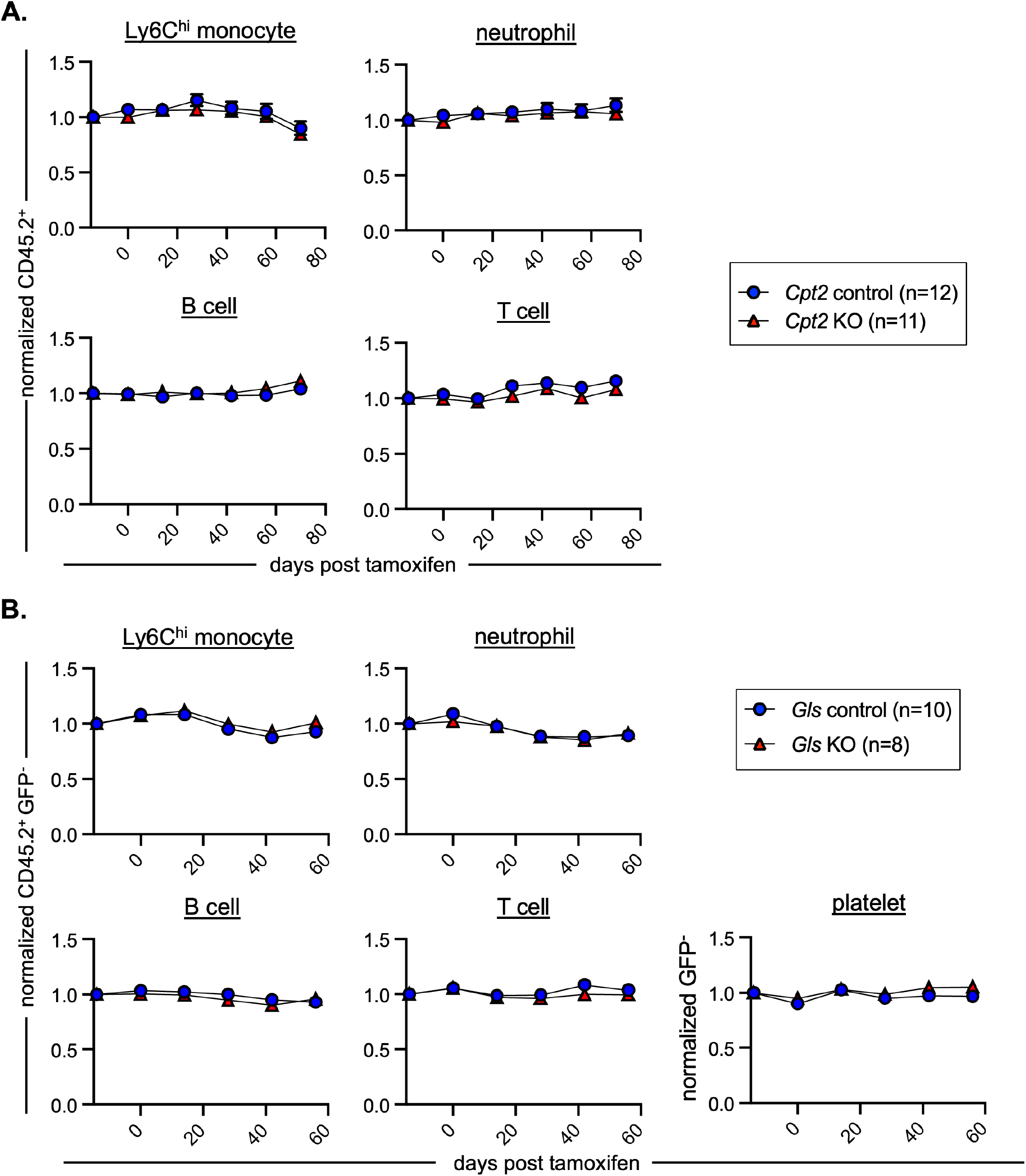
related to Figure 3. Quantification of peripheral blood chimerism of CPT2*-* and GLS-deficient chimeras. (A-B) CD45.2 peripheral blood chimerism of mature cell populations was assessed every 2 weeks in *Cpt2* (A) or *Gls* (B) chimeras. Values are normalized to pre-tamoxifen chimerism of each cell type. Mean values ± SEM are shown.

**Figure S4.**
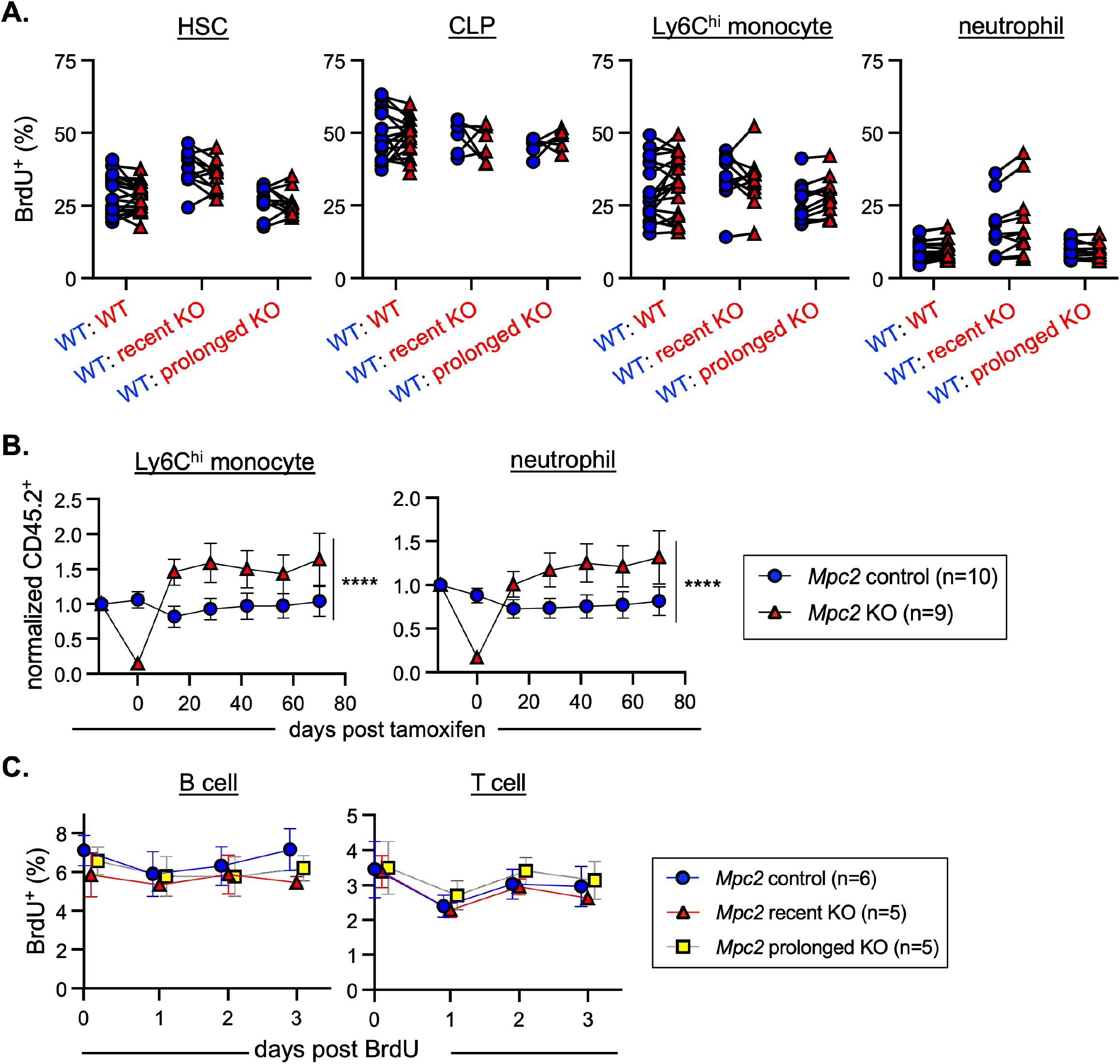
related to Figure 4. Proliferation, survival, and recovery of MPC2- deficient cells. (A) BrdU incorporation was measured in both CD45.1^+^ (wild-type) and CD45.2^+^ (wild-type, recent *Mpc2* deletion, or prolonged *Mpc2* deletion) HSCs, CLPs, Ly6C^hi^ monocytes, and neutrophils following a 1-hour pulse of BrdU. Each line connects CD45.1^+^ and CD45.2^+^ cells within the same mouse. Data are pooled from two independent experiments. (B) *Mpc2* chimeras were setup at a ratio of 90% wild-type to 10% *Mpc2^fl/fl^; ROSA26 CreER^-/-^* or *^+/-^* bone marrow cells (as opposed to the 1:1 ratio in Figure 3). CD45.2 peripheral blood chimerism of Ly6C^hi^ monocytes and neutrophils was assessed every 2 weeks in *Mpc2* chimeras. Values are normalized to pre-tamoxifen chimerism of each cell type. Mean values ± SEM are shown. ****p<0.0001 by paired 2-way ANOVA with post-hoc Tukey’s multiple comparisons test. (C) BrdU incorporation was measured in peripheral CD45.2^+^ B and T cells for 3 days following a week of BrdU water administration. Mean values ± SEM are shown.

**Figure S5.**
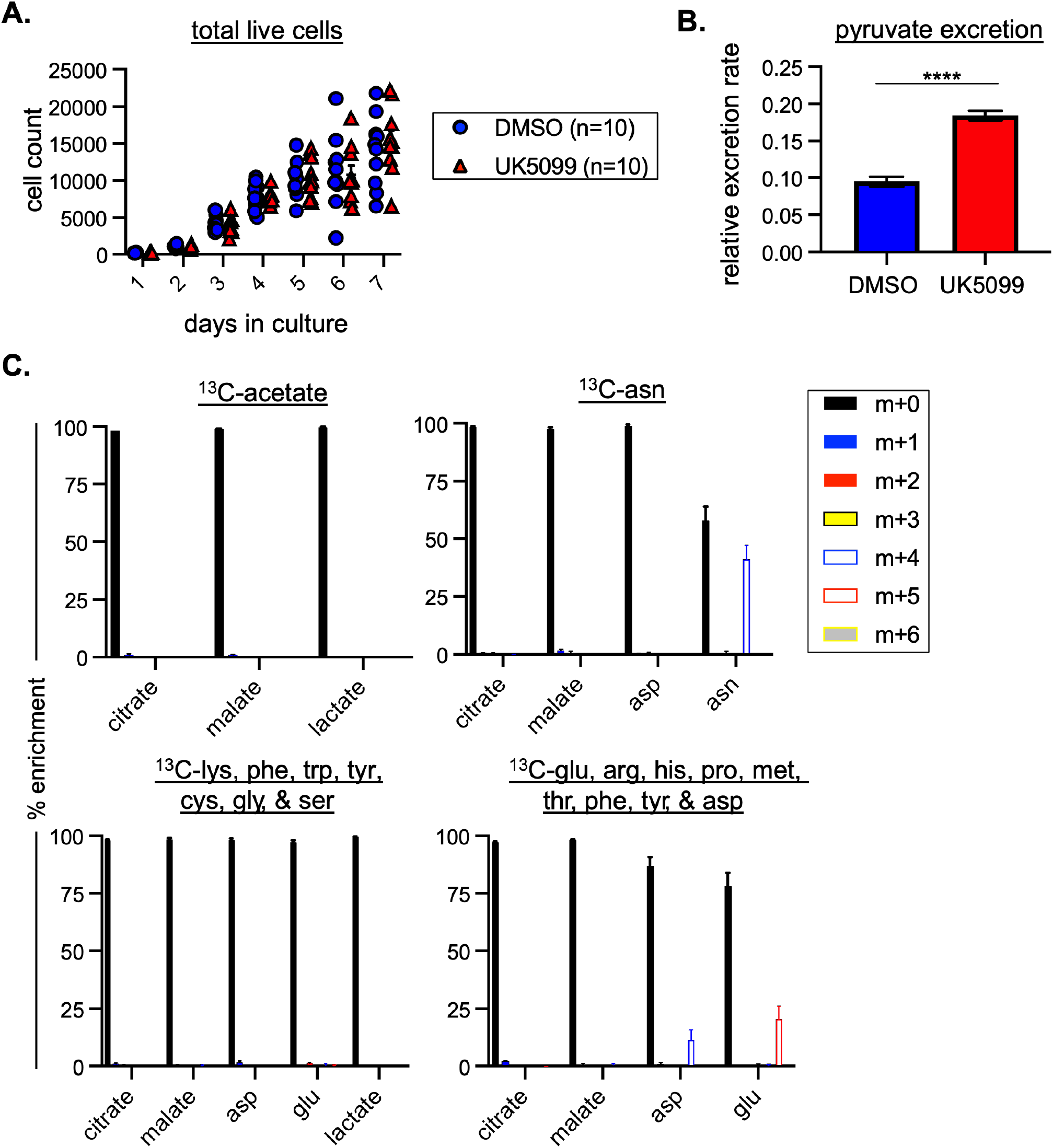
related to Figure 5. Generation and usage of pyruvate by GMPs *in vitro*. (A) Wild-type GMPs were cultured with DMSO or 10μM UK5099 for 7 days. On each day of the culture, cells were harvested, and live cells were counted. One experiment is depicted as a representative of four independent experiments. (B) Pyruvate excretion rate in the media of GMPs cultured with DMSO or UK5099 over a 48-hour culture. ****p<0.0001 by Student’s 2-tailed t-test. Mean values ± SEM are shown for 4 replicates. (C) LC/MS analysis of ^13^C incorporation into TCA cycle intermediates following a 24-hour culture of wild-type GMPs with uniformly ^13^C-labeled acetate, asparagine, lysine/phenylalanine/tryptophan/tyrosine/cysteine/glycine/serine, or glutamate/arginine/proline/methionine/threonine/phenylalanine/tyrosine/aspartate. Labeling data were corrected for natural-abundance ^13^C. Mean values ± SEM are shown for 3-5 replicates.

**Figure S6.**
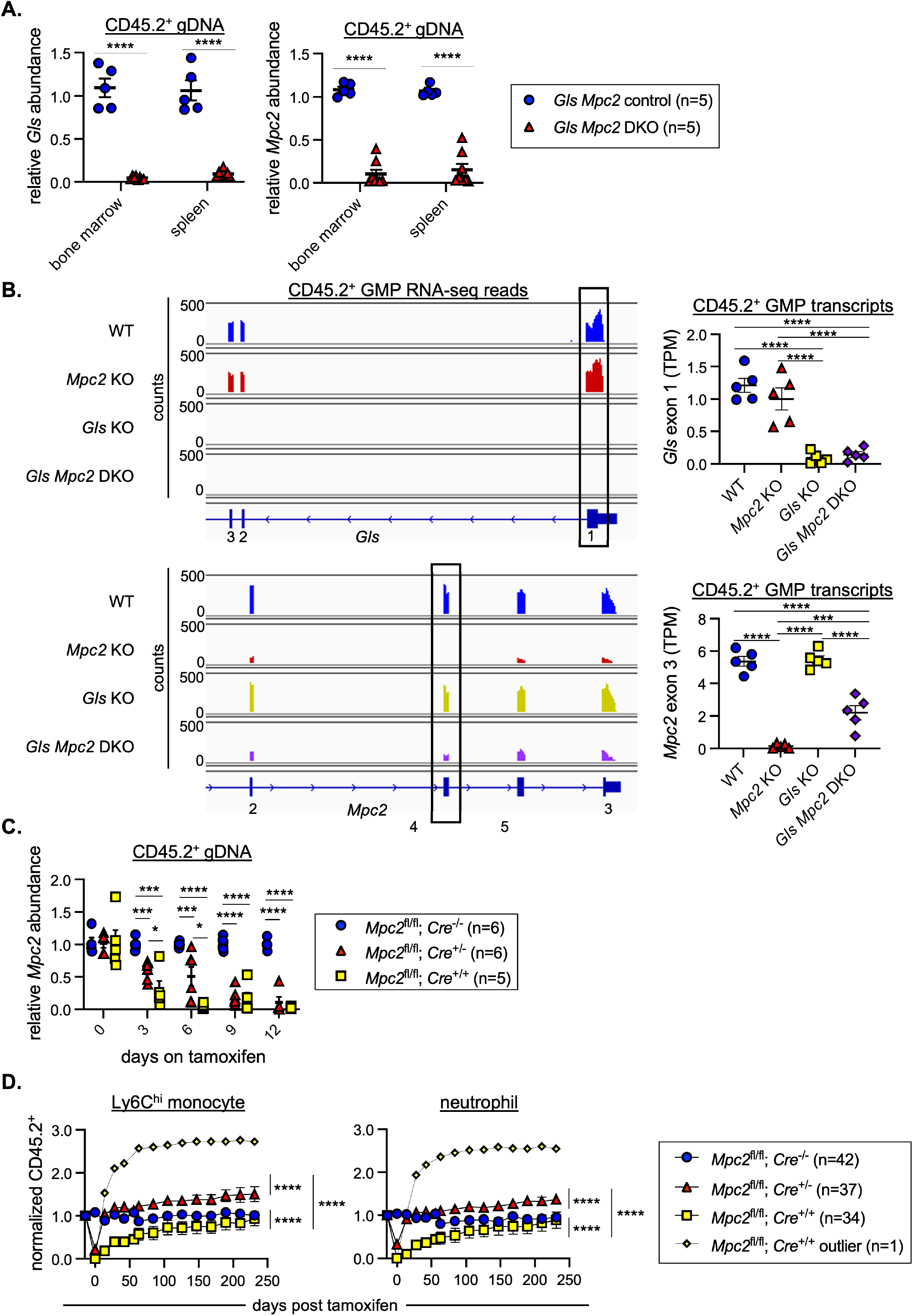
related to Figure 6. Quantification of efficiency and rates of *Gls* and *Mpc2* deletion. (A) Quantitative PCR analysis of genomic *Gls* and *Mpc2* deletion in recent *Gls Mpc2* KO chimeras. CD45.2^+^ cells from the bone marrow or spleen were sorted for genomic DNA extraction. DNA quantification within loxP sites was quantified relative to wild-type cells and GAPDH. Mean values ± SEM are shown. ****p<0.0001 by Student’s 2-tailed t-test. (B) RNA-seq reads across exons 1-3 of *Gls* (top) and 2-5 of *Mpc2* (bottom) of CD45.2^+^ GMPs from *ROSA26 CreER^+/-^* (WT), *Mpc2* KO, *Gls* KO, or *Gls Mpc2* DKO chimeras about 10 weeks after tamoxifen treatment. Exon 1 of *Gls* and exon 3 of *Mpc2* are floxed in the appropriate genotyped mice. One representative trace for each genotype is shown using Integrative Genomics Viewer (IGV). Transcripts per kilobase million (TPMs) were quantified for the floxed exon of *Gls* (top) or *Mpc2* (bottom). TPMs were calculated from reads obtained using DEXSeq. Mean values ± SEM are shown. ***p<0.001 and ****p<0.0001 by 1-way ANOVA with post-hoc Tukey’s multiple comparisons test. (C) Quantitative PCR analysis of genomic *Mpc2* deletion in *Mpc2* chimeras. *Mpc2* chimeras were setup with *Mpc2^fl/fl^; ROSA26 CreER^-/-^*, *^+/-^*, or *^+/+^* bone marrow cells. CD45.2^+^ cells from the blood were sorted for genomic DNA extraction on days 0, 3, 6, 9, and 12 of tamoxifen administration. DNA quantification within loxP sites was quantified relative to wild-type cells and GAPDH. Mean values ± SEM are shown. Data are pooled from two independent experiments. *p<0.05, ***p<0.001, and ****p<0.0001 by 1-way ANOVA with post-hoc Tukey’s multiple comparisons test. (D) *Mpc2* chimeras were setup with *Mpc2^fl/fl^; ROSA26 CreER^-/-^*, *^+/-^*, or *^+/+^* bone marrow cells. CD45.2 peripheral blood chimerism of Ly6C^hi^ monocytes and neutrophils was assessed every 2-3 weeks. Values are normalized to pre-tamoxifen chimerism of each cell type. *Mpc2^fl/fl^; ROSA26 CreER^-/-^* and ^+/-^ data is reused from Figure 3. One *Mpc2^fl/fl^; ROSA26 CreER^+/+^* recipient chimera was found to be a significant outlier and was separated and omitted from statistical analysis of that group. Data are pooled from three independent experiments. ****p<0.0001 by paired 2-way ANOVA with post-hoc Tukey’s multiple comparisons test.

## References

Agathocleous, M., Meacham, C. E., Burgess, R. J., Piskounova, E., Zhao, Z., Crane, G. M., Cowin, B. L., Bruner, E., Murphy, M. M., Chen, W., Spangrude, G. J., Hu, Z., DeBerardinis, R. J., & Morrison, S. J. (2017). Ascorbate regulates haematopoietic stem cell function and leukaemogenesis. Nature, 549(7673), 476–481. https://doi.org/10.1038/nature23876

Anders, S., Reyes, A., & Huber, W. (2012). Detecting differential usage of exons from RNA-seq data. Genome Research, 22(10), 2008–2017. https://doi.org/10.1101/gr.133744.111

Ansó, E., Weinberg, S. E., Diebold, L. P., Thompson, B. J., Malinge, S., Schumacker, P. T., Liu, X., Zhang, Y., Shao, Z., Steadman, M., Marsh, K. M., Xu, J., Crispino, J. D., & Chandel, N. S. (2017). The mitochondrial respiratory chain is essential for haematopoietic stem cell function. Nature Cell Biology, 19(6), 614–625. https://doi.org/10.1038/ncb3529

Arison, R. N., Ciaccio, E. I., Glitzer, M. S., Cassaro, J. A., & Pruss, M. P. (1967). Light and electron microscopy of lesions in rats rendered diabetic with streptozotocin. Diabetes, 16(1), 51–56. https://doi.org/10.2337/diab.16.1.51

Bejarano-García, J. A., Millán-Uclés, Á., Rosado, I. V., Sánchez-Abarca, L. I., Caballero- Velázquez, T., Durán-Galván, M. J., Pérez-Simón, J. A., & Piruat, J. I. (2016). Sensitivity of hematopoietic stem cells to mitochondrial dysfunction by SdhD gene deletion. Cell Death & Disease, 7(12), e2516–e2516. https://doi.org/10.1038/cddis.2016.411

Bensard, C. L., Wisidagama, D. R., Olson, K. A., Berg, J. A., Krah, N. M., Schell, J. C., Nowinski, S. M., Fogarty, S., Bott, A. J., Wei, P., Dove, K. K., Tanner, J. M., Panic, V., Cluntun, A., Lettlova, S., Earl, C. S., Namnath, D. F., Vázquez-Arreguín, K., Villanueva, C. J., … Rutter, J. (2020). Regulation of Tumor Initiation by the Mitochondrial Pyruvate Carrier. Cell Metabolism, 31(2), 284–300.e7. https://doi.org/10.1016/j.cmet.2019.11.002

Bhattacharya, D., Rossi, D. J., Bryder, D., & Weissman, I. L. (2006). Purified hematopoietic stem cell engraftment of rare niches corrects severe lymphoid deficiencies without host conditioning. The Journal of Experimental Medicine, 203(1), 73–85. https://doi.org/10.1084/jem.20051714

Boettcher, S., & Manz, M. G. (2017). Regulation of Inflammation- and Infection-Driven Hematopoiesis. Trends in Immunology, 38(5), 345–357. https://doi.org/10.1016/j.it.2017.01.004

Born, J., Lange, T., Hansen, K., Mölle, M., & Fehm, H. L. (1997). Effects of sleep and circadian rhythm on human circulating immune cells. The Journal of Immunology, 158, 4454–4464.

Bricker, D. K., Taylor, E. B., Schell, J. C., Orsak, T., Boutron, A., Chen, Y.-C., Cox, J. E., Cardon, C. M., Van Vranken, J. G., Dephoure, N., Redin, C., Boudina, S., Gygi, S. P., Brivet, M., Thummel, C. S., & Rutter, J. (2012). A mitochondrial pyruvate carrier required for pyruvate uptake in yeast, Drosophila, and humans. Science (New York, N.Y.), 337(6090), 96–100. https://doi.org/10.1126/science.1218099

Bryder, D., Rossi, D. J., & Weissman, I. L. (2006). Hematopoietic Stem Cells. The American Journal of Pathology, 169(2), 338–346. https://doi.org/10.2353/ajpath.2006.060312

Chen, C., Liu, Y., Liu, R., Ikenoue, T., Guan, K.-L., Liu, Y., & Zheng, P. (2008). TSC– mTOR maintains quiescence and function of hematopoietic stem cells by repressing mitochondrial biogenesis and reactive oxygen species. Journal of Experimental Medicine, 205(10), 2397–2408. https://doi.org/10.1084/jem.20081297

Cheng, T., Sudderth, J., Yang, C., Mullen, A. R., Jin, E. S., Matés, J. M., & DeBerardinis, R. J. (2011). Pyruvate carboxylase is required for glutamine-independent growth of tumor cells. Proceedings of the National Academy of Sciences of the United States of America, 108(21), 8674–8679. https://doi.org/10.1073/pnas.1016627108

Cheshier, S. H., Prohaska, S. S., & Weissman, I. L. (2007). The effect of bleeding on hematopoietic stem cell cycling and self-renewal. Stem Cells and Development, 16(5), 707–717. https://doi.org/10.1089/scd.2007.0017

Cronkite, E. P., Fliedner, T. M., Bond, V. P., Rubini, J. R., Brecher, G., & Quastler, H. (1959). Dynamics of Hemopoietic Proliferation in Man and Mice Studied by H3- Thymidine Incorporation into Dna*. Annals of the New York Academy of Sciences, 77(3), 803–820. https://doi.org/10.1111/j.1749-6632.1959.tb36943.x

Curthoys, N. P., & Watford, M. (1995). Regulation of glutaminase activity and glutamine metabolism. Annual Review of Nutrition, 15, 133–159. https://doi.org/10.1146/annurev.nu.15.070195.001025

Davidson, S. M., Papagiannakopoulos, T., Olenchock, B. A., Heyman, J. E., Keibler, M. A., Luengo, A., Bauer, M. R., Jha, A. K., O’Brien, J. P., Pierce, K. A., Gui, D. Y., Sullivan, L. B., Wasylenko, T. M., Subbaraj, L., Chin, C. R., Stephanopolous, G., Mott, B. T., Jacks, T., Clish, C. B., & Vander Heiden, M. G. (2016). Environment Impacts the Metabolic Dependencies of Ras-Driven Non-Small Cell Lung Cancer. Cell Metabolism, 23(3), 517–528. https://doi.org/10.1016/j.cmet.2016.01.007

DeBerardinis, R. J., Lum, J. J., Hatzivassiliou, G., & Thompson, C. B. (2008). The Biology of Cancer: Metabolic Reprogramming Fuels Cell Growth and Proliferation. Cell Metabolism, 7(1), 11–20. https://doi.org/10.1016/j.cmet.2007.10.002

DeBerardinis, R. J., Mancuso, A., Daikhin, E., Nissim, I., Yudkoff, M., Wehrli, S., & Thompson, C. B. (2007). Beyond aerobic glycolysis: Transformed cells can engage in glutamine metabolism that exceeds the requirement for protein and nucleotide synthesis. Proceedings of the National Academy of Sciences, 104(49), 19345–19350. https://doi.org/10.1073/pnas.0709747104

Faubert, B., & DeBerardinis, R. J. (2017). Analyzing Tumor Metabolism In Vivo. Annual Review of Cancer Biology, 1(1), 99–117. https://doi.org/10.1146/annurev-cancerbio-050216-121954

Forsberg, E. C., Serwold, T., Kogan, S., Weissman, I. L., & Passegué, E. (2006). New evidence supporting megakaryocyte-erythrocyte potential of flk2/flt3+ multipotent hematopoietic progenitors. Cell, 126(2), 415–426. https://doi.org/10.1016/j.cell.2006.06.037

Fulcher, D. A., & Basten, A. (1997). B cell life span: A review. Immunology & Cell Biology, 75(5), 446–455. https://doi.org/10.1038/icb.1997.69

Gan, B., Hu, J., Jiang, S., Liu, Y., Sahin, E., Zhuang, L., Fletcher-Sananikone, E., Colla, S., Wang, Y. A., Chin, L., & DePinho, R. A. (2010). Lkb1 regulates quiescence and metabolic homeostasis of haematopoietic stem cells. Nature, 468(7324), 701–704. https://doi.org/10.1038/nature09595

Garland, P. B., Randle, P. J., & Newsholme, E. A. (1963). Citrate as an Intermediary in the Inhibition of Phosphofructokinase in Rat Heart Muscle by Fatty Acids, Ketone Bodies, Pyruvate, Diabetes and Starvation. Nature, 200(4902), 169–170. https://doi.org/10.1038/200169a0

Guitart, A. V., Panagopoulou, T. I., Villacreces, A., Vukovic, M., Sepulveda, C., Allen, L., Carter, R. N., van de Lagemaat, L. N., Morgan, M., Giles, P., Sas, Z., Gonzalez, M. V., Lawson, H., Paris, J., Edwards-Hicks, J., Schaak, K., Subramani, C., Gezer, D., Armesilla-Diaz, A., … Kranc, K. R. (2017). Fumarate hydratase is a critical metabolic regulator of hematopoietic stem cell functions. Journal of Experimental Medicine, 214(3), 719–735. https://doi.org/10.1084/jem.20161087

Gurumurthy, S., Xie, S. Z., Alagesan, B., Kim, J., Yusuf, R. Z., Saez, B., Tzatsos, A., Ozsolak, F., Milos, P., Ferrari, F., Park, P. J., Shirihai, O. S., Scadden, D. T., & Bardeesy, N. (2010). The Lkb1 metabolic sensor maintains haematopoietic stem cell survival. Nature, 468(7324), 659–663. https://doi.org/10.1038/nature09572

Halestrap, A. P., & Denton, R. M. (1975). The specificity and metabolic implications of the inhibition of pyruvate transport in isolated mitochondria and intact tissue preparations by alpha-Cyano-4-hydroxycinnamate and related compounds. The Biochemical Journal, 148(1), 97–106. https://doi.org/10.1042/bj1480097

Hattori, A., Tsunoda, M., Konuma, T., Kobayashi, M., Nagy, T., Glushka, J., Tayyari, F., McSkimming, D., Kannan, N., Tojo, A., Edison, A. S., & Ito, T. (2017). Cancer progression by reprogrammed BCAA metabolism in myeloid leukaemia. Nature, 545(7655), 500–504. https://doi.org/10.1038/nature22314

Haus, E., Lakatua, D. J., Swoyer, J., & Sackett-Lundeen, L. (1983). Chronobiology in hematology and immunology. American Journal of Anatomy, 168(4), 467–517. https://doi.org/10.1002/aja.1001680406

Heng, T. S., Painter, M. W., & Project, U. I. G. (2008). The Immunological Genome Project: Networks of gene expression in immune cells. Nature Immunology, 9(10), 1091–1094.

Herzig, S., Raemy, E., Montessuit, S., Veuthey, J.-L., Zamboni, N., Westermann, B., Kunji, E. R. S., & Martinou, J.-C. (2012). Identification and functional expression of the mitochondrial pyruvate carrier. Science (New York, N.Y.), 337(6090), 93–96. https://doi.org/10.1126/science.1218530

Hinge, A., He, J., Bartram, J., Javier, J., Xu, J., Fjellman, E., Sesaki, H., Li, T., Yu, J., Wunderlich, M., Mulloy, J., Kofron, M., Salomonis, N., Grimes, H. L., & Filippi, M.-D. (2020). Asymmetrically Segregated Mitochondria Provide Cellular Memory of Hematopoietic Stem Cell Replicative History and Drive HSC Attrition. Cell Stem Cell, 26(3), 420–430.e6. https://doi.org/10.1016/j.stem.2020.01.016

Ho, T. T., Warr, M. R., Adelman, E. R., Lansinger, O. M., Flach, J., Verovskaya, E. V., Figueroa, M. E., & Passegué, E. (2017). Autophagy maintains the metabolism and function of young and old (hematopoietic) stem cells. Nature, 543(7644), 205–210. https://doi.org/10.1038/nature21388

Houten, S. M., Violante, S., Ventura, F. V., & Wanders, R. J. A. (2016). The Biochemistry and Physiology of Mitochondrial Fatty Acid β-Oxidation and Its Genetic Disorders. Annual Review of Physiology, 78(1), 23–44. https://doi.org/10.1146/annurev-physiol-021115-105045

Hutson, S. M., Fenstermacher, D., & Mahar, C. (1988). Role of mitochondrial transamination in branched chain amino acid metabolism. Journal of Biological Chemistry, 263(8), 3618–3625. https://doi.org/10.1016/S0021-9258(18)68969-0

Ichihara, A., & Koyama, E. (1966). Transaminase of Branched Chain Amino Acids. The Journal of Biochemistry, 59(2), 160–169.

Inoue, S.-I., Noda, S., Kashima, K., Nakada, K., Hayashi, J.-I., & Miyoshi, H. (2010). Mitochondrial respiration defects modulate differentiation but not proliferation of hematopoietic stem and progenitor cells. FEBS Letters, 584(15), 3402–3409. https://doi.org/10.1016/j.febslet.2010.06.036

Isackson, P. J., Bennett, M. J., Lichter-Konecki, U., Willis, M., Nyhan, W. L., Sutton, V. R., Tein, I., & Vladutiu, G. D. (2008). CPT2 gene mutations resulting in lethal neonatal or severe infantile carnitine palmitoyltransferase II deficiency. Molecular Genetics and Metabolism, 94(4), 422–427. https://doi.org/10.1016/j.ymgme.2008.05.002

Ito, K., Bonora, M., & Ito, K. (2019). Metabolism as master of hematopoietic stem cell fate. International Journal of Hematology, 109(1), 18–27. https://doi.org/10.1007/s12185-018-2534-z

Ito, K., Hirao, A., Arai, F., Matsuoka, S., Takubo, K., Hamaguchi, I., Nomiyama, K., Hosokawa, K., Sakurada, K., Nakagata, N., Ikeda, Y., Mak, T. W., & Suda, T. (2004). Regulation of oxidative stress by ATM is required for self-renewal of haematopoietic stem cells. Nature, 431(7011), 997–1002. https://doi.org/10.1038/nature02989

Ito, K., Hirao, A., Arai, F., Takubo, K., Matsuoka, S., Miyamoto, K., Ohmura, M., Naka, K., Hosokawa, K., Ikeda, Y., & Suda, T. (2006). Reactive oxygen species act through p38 MAPK to limit the lifespan of hematopoietic stem cells. Nature Medicine, 12(4), 446–451. https://doi.org/10.1038/nm1388

Ito, K., & Suda, T. (2014). Metabolic requirements for the maintenance of self-renewing stem cells. Nature Reviews. Molecular Cell Biology, 15(4), 243–256. https://doi.org/10.1038/nrm3772

Ji, S., You, Y., Kerner, J., Hoppel, C. L., Schoeb, T. R., Chick, W. S. H., Hamm, D. A., Sharer, J. D., & Wood, P. A. (2008). Homozygous carnitine palmitoyltransferase 1b (muscle isoform) deficiency is lethal in the mouse. Molecular Genetics and Metabolism, 93(3), 314–322. https://doi.org/10.1016/j.ymgme.2007.10.006

Jun, S., Mahesula, S., Mathews, T. P., Martin-Sandoval, M. S., Zhao, Z., Piskounova, E., & Agathocleous, M. (2021). The requirement for pyruvate dehydrogenase in leukemogenesis depends on cell lineage. Cell Metabolism, 33(9), 1777–1792.e8. https://doi.org/10.1016/j.cmet.2021.07.016

Kim, D., Langmead, B., & Salzberg, S. L. (2015). HISAT: A fast spliced aligner with low memory requirements. Nature Methods, 12(4), 357–360. https://doi.org/10.1038/nmeth.3317

Lam, W. Y., Becker, A. M., Kennerly, K. M., Wong, R., Curtis, J. D., Llufrio, E. M., McCommis, K. S., Fahrmann, J., Pizzato, H. A., Nunley, R. M., Lee, J., Wolfgang, M. J., Patti, G. J., Finck, B. N., Pearce, E. L., & Bhattacharya, D. (2016). Mitochondrial Pyruvate Import Promotes Long-Term Survival of Antibody-Secreting Plasma Cells. Immunity, 45(1), 60–73. https://doi.org/10.1016/j.immuni.2016.06.011

Lawrence, J. S., Stephens, D. J., & Jones, E. (1933). Studies in the Normal Human White Blood Cell Picture. II. The Effect of Digestion on the White Blood Cells. American Journal of Physiology-Legacy Content. https://doi.org/10.1152/ajplegacy.1933.106.2.309

Lee, J., Ellis, J. M., & Wolfgang, M. J. (2015). Adipose Fatty Acid Oxidation Is Required for Thermogenesis and Potentiates Oxidative Stress-Induced Inflammation. Cell Reports, 10(2), 266–279. https://doi.org/10.1016/j.celrep.2014.12.023

Lenzen, S. (2008). The mechanisms of alloxan- and streptozotocin-induced diabetes. Diabetologia, 51(2), 216–226. https://doi.org/10.1007/s00125-007-0886-7

Longo, N., Amat di San Filippo, C., & Pasquali, M. (2006). Disorders of carnitine transport and the carnitine cycle. American Journal of Medical Genetics. Part C, Seminars in Medical Genetics, 142C(2), 77–85. https://doi.org/10.1002/ajmg.c.30087

Love, M. I., Huber, W., & Anders, S. (2014). Moderated estimation of fold change and dispersion for RNA-seq data with DESeq2. Genome Biology, 15(12), 550. https://doi.org/10.1186/s13059-014-0550-8

Luchsinger, L. L., de Almeida, M. J., Corrigan, D. J., Mumau, M., & Snoeck, H.-W. (2016). Mitofusin 2 maintains haematopoietic stem cells with extensive lymphoid potential. Nature, 529(7587), 528–531. https://doi.org/10.1038/nature16500

Luo, Y., Chen, G.-L., Hannemann, N., Ipseiz, N., Krönke, G., Bäuerle, T., Munos, L., Wirtz, S., Schett, G., & Bozec, A. (2015). Microbiota from Obese Mice Regulate Hematopoietic Stem Cell Differentiation by Altering the Bone Niche. Cell Metabolism, 22(5), 886–894. https://doi.org/10.1016/j.cmet.2015.08.020

Mansell, E., Sigurdsson, V., Deltcheva, E., Brown, J., James, C., Miharada, K., Soneji, S., Larsson, J., & Enver, T. (2021). Mitochondrial Potentiation Ameliorates Age-Related Heterogeneity in Hematopoietic Stem Cell Function. Cell Stem Cell, 28(2), 241–256.e6. https://doi.org/10.1016/j.stem.2020.09.018

Maryanovich, M., Oberkovitz, G., Niv, H., Vorobiyov, L., Zaltsman, Y., Brenner, O., Lapidot, T., Jung, S., & Gross, A. (2012). The ATM–BID pathway regulates quiescence and survival of haematopoietic stem cells. Nature Cell Biology, 14(5), 535–541. https://doi.org/10.1038/ncb2468

Maryanovich, M., Zaltsman, Y., Ruggiero, A., Goldman, A., Shachnai, L., Zaidman, S. L., Porat, Z., Golan, K., Lapidot, T., & Gross, A. (2015). An MTCH2 pathway repressing mitochondria metabolism regulates haematopoietic stem cell fate. Nature Communications, 6, 7901. https://doi.org/10.1038/ncomms8901

Masson, J., Darmon, M., Conjard, A., Chuhma, N., Ropert, N., Thoby-Brisson, M., Foutz, A. S., Parrot, S., Miller, G. M., Jorisch, R., Polan, J., Hamon, M., Hen, R., & Rayport, S. (2006). Mice Lacking Brain/Kidney Phosphate-Activated Glutaminase Have Impaired Glutamatergic Synaptic Transmission, Altered Breathing, Disorganized Goal-Directed Behavior and Die Shortly after Birth. The Journal of Neuroscience, 26(17), 4660–4671. https://doi.org/10.1523/JNEUROSCI.4241-05.2006

Mauer, A. M., Athens, J. W., Ashenbrucker, H., Cartwright, G. E., & Wintrobe, M. M. (1960). Leukokinetic Studies. II. A Method for Labeling Granulocytes in Vitro with Radioactive Diisopropylfluorophosphate (DFP). The Journal of Clinical Investigation, 39(9), 1481–1486. https://doi.org/10.1172/JCI104167

McCommis, K. S., Chen, Z., Fu, X., McDonald, W. G., Colca, J. R., Kletzien, R. F., Burgess, S. C., & Finck, B. N. (2015). Loss of Mitochondrial Pyruvate Carrier 2 in Liver Leads to Defects in Gluconeogenesis and Compensation via Pyruvate-Alanine Cycling. Cell Metabolism, 22(4), 682–694. https://doi.org/10.1016/j.cmet.2015.07.028

Mingote, S., Masson, J., Gellman, C., Thomsen, G. M., Lin, C.-S., Merker, R. J., Gaisler-Salomon, I., Wang, Y., Ernst, R., Hen, R., & Rayport, S. (2016). Genetic Pharmacotherapy as an Early CNS Drug Development Strategy: Testing Glutaminase Inhibition for Schizophrenia Treatment in Adult Mice. Frontiers in Systems Neuroscience, 9. https://doi.org/10.3389/fnsys.2015.00165

Mortensen, M., Soilleux, E. J., Djordjevic, G., Tripp, R., Lutteropp, M., Sadighi-Akha, E., Stranks, A. J., Glanville, J., Knight, S., W. Jacobsen, S.-E., Kranc, K. R., & Simon, A. K. (2011). The autophagy protein Atg7 is essential for hematopoietic stem cell maintenance. Journal of Experimental Medicine, 208(3), 455–467. https://doi.org/10.1084/jem.20101145

Muller, Y. D., Golshayan, D., Ehirchiou, D., Wyss, J. C., Giovannoni, L., Meier, R., Serre-Beinier, V., Puga Yung, G., Morel, P., Bühler, L. H., & Seebach, J. D. (2011). Immunosuppressive Effects of Streptozotocin-Induced Diabetes Result in Absolute Lymphopenia and a Relative Increase of T Regulatory Cells. Diabetes, 60(9), 2331–2340. https://doi.org/10.2337/db11-0159

Nagareddy, P. R., Murphy, A. J., Stirzaker, R. A., Hu, Y., Yu, S., Miller, R. G., Ramkhelawon, B., Distel, E., Westerterp, M., Huang, L.-S., Schmidt, A. M., Orchard, T. J., Fisher, E. A., Tall, A. R., & Goldberg, I. J. (2013). Hyperglycemia promotes myelopoiesis and impairs the resolution of atherosclerosis. Cell Metabolism, 17(5), 695–708. https://doi.org/10.1016/j.cmet.2013.04.001

Nakada, D., Saunders, T. L., & Morrison, S. J. (2010). Lkb1 regulates cell cycle and energy metabolism in haematopoietic stem cells. Nature, 468(7324), 653–658. https://doi.org/10.1038/nature09571

Nakamura-Ishizu, A., Ito, K., & Suda, T. (2020). Hematopoietic Stem Cell Metabolism during Development and Aging. Developmental Cell, 54(2), 239–255. https://doi.org/10.1016/j.devcel.2020.06.029

Norddahl, G. L., Pronk, C. J., Wahlestedt, M., Sten, G., Nygren, J. M., Ugale, A., Sigvardsson, M., & Bryder, D. (2011). Accumulating mitochondrial DNA mutations drive premature hematopoietic aging phenotypes distinct from physiological stem cell aging. Cell Stem Cell, 8(5), 499–510. https://doi.org/10.1016/j.stem.2011.03.009

Nyman, L. R., Cox, K. B., Hoppel, C. L., Kerner, J., Barnoski, B. L., Hamm, D. A., Tian, L., Schoeb, T. R., & Wood, P. A. (2005). Homozygous carnitine palmitoyltransferase 1a (liver isoform) deficiency is lethal in the mouse. Molecular Genetics and Metabolism, 86(1–2), 179–187. https://doi.org/10.1016/j.ymgme.2005.07.021

Oburoglu, L., Tardito, S., Fritz, V., de Barros, S. C., Merida, P., Craveiro, M., Mamede, J., Cretenet, G., Mongellaz, C., An, X., Klysz, D., Touhami, J., Boyer-Clavel, M., Battini, J.-L., Dardalhon, V., Zimmermann, V. S., Mohandas, N., Gottlieb, E., Sitbon, M., … Taylor, N. (2014). Glucose and Glutamine Metabolism Regulate Human Hematopoietic Stem Cell Lineage Specification. Cell Stem Cell, 15(2), 169–184. https://doi.org/10.1016/j.stem.2014.06.002

Passegué, E., Wagers, A. J., Giuriato, S., Anderson, W. C., & Weissman, I. L. (2005). Global analysis of proliferation and cell cycle gene expression in the regulation of hematopoietic stem and progenitor cell fates. The Journal of Experimental Medicine, 202(11), 1599–1611. https://doi.org/10.1084/jem.20050967

Patro, R., Duggal, G., Love, M. I., Irizarry, R. A., & Kingsford, C. (2017). Salmon provides fast and bias-aware quantification of transcript expression. Nature Methods, 14(4), 417–419. https://doi.org/10.1038/nmeth.4197

Pickel, L., & Sung, H.-K. (2020). Feeding Rhythms and the Circadian Regulation of Metabolism. Frontiers in Nutrition, 7, 39. https://doi.org/10.3389/fnut.2020.00039

Pillay, J., den Braber, I., Vrisekoop, N., Kwast, L. M., de Boer, R. J., Borghans, J. A. M., Tesselaar, K., & Koenderman, L. (2010). In vivo labeling with 2H2O reveals a human neutrophil lifespan of 5.4 days. Blood, 116(4), 625–627. https://doi.org/10.1182/blood-2010-01-259028

Poorman, R. A., Randolph, A., Kemp, R. G., & Heinrikson, R. L. (1984). Evolution of phosphofructokinase—Gene duplication and creation of new effector sites. Nature, 309(5967), 467–469. https://doi.org/10.1038/309467a0

Potter, G. D. M., Cade, J. E., Grant, P. J., & Hardie, L. J. (2016). Nutrition and the Circadian System. The British Journal of Nutrition, 116(3), 434–442. https://doi.org/10.1017/S0007114516002117

Qi, L., Martin-Sandoval, M. S., Merchant, S., Gu, W., Eckhardt, M., Mathews, T. P., Zhao, Z., Agathocleous, M., & Morrison, S. J. (2021). Aspartate availability limits hematopoietic stem cell function during hematopoietic regeneration. Cell Stem Cell. https://doi.org/10.1016/j.stem.2021.07.011

Raffel, S., Falcone, M., Kneisel, N., Hansson, J., Wang, W., Lutz, C., Bullinger, L., Poschet, G., Nonnenmacher, Y., Barnert, A., Bahr, C., Zeisberger, P., Przybylla, A., Sohn, M., Tönjes, M., Erez, A., Adler, L., Jensen, P., Scholl, C., … Trumpp, A. (2017). BCAT1 restricts αKG levels in AML stem cells leading to IDHmut-like DNA hypermethylation. Nature, 551(7680), 384–388. https://doi.org/10.1038/nature24294

Rakieten, N., Rakieten, M. L., & Nadkarni, M. V. (1963). Studies on the diabetogenic action of streptozotocin (NSC-37917). Cancer Chemotherapy Reports, 29, 91–98.

Ramstead, A. G., Wallace, J. A., Lee, S.-H., Bauer, K. M., Tang, W. W., Ekiz, H. A., Lane, T. E., Cluntun, A. A., Bettini, M. L., Round, J. L., Rutter, J., & O’Connell, R. M. (2020). Mitochondrial Pyruvate Carrier 1 Promotes Peripheral T Cell Homeostasis through Metabolic Regulation of Thymic Development. Cell Reports, 30(9), 2889–2899.e6. https://doi.org/10.1016/j.celrep.2020.02.042

Sabin, F., Cunningham, R., Doan, C., & Kindwale, J. (1927). The normal rhythm of white blood cells. Bull. Johns Hopkins Hosp, 37, 14–67.

Schaffer, J. E., & Lodish, H. F. (1994). Expression cloning and characterization of a novel adipocyte long chain fatty acid transport protein. Cell, 79(3), 427–436. https://doi.org/0092-8674(94)90252-6 [pii]

Schell, J. C., Olson, K. A., Jiang, L., Hawkins, A. J., VanVranken, J. G., Xie, J., Egnatchik, R. A., Earl, E. G., DeBerardinis, R. J., & Rutter, J. (2014). A role for the mitochondrial pyruvate carrier as a repressor of the warburg effect and colon cancer cell growth. Molecular Cell, 56(3), 400–413. https://doi.org/10.1016/j.molcel.2014.09.026

Schlierf, G., & Dorow, E. (1973). Diurnal Patterns of Triglycerides, Free Fatty Acids, Blood Sugar, and Insulin during Carbohydrate-Induction in Man and Their Modification by Nocturnal Suppression of Lipolysis. The Journal of Clinical Investigation, 52(3), 732–740. https://doi.org/10.1172/JCI107235

Sellers, K., Fox, M. P., Bousamra, M., Slone, S. P., Higashi, R. M., Miller, D. M., Wang, Y., Yan, J., Yuneva, M. O., Deshpande, R., Lane, A. N., & Fan, T. W.-M. (2015). Pyruvate carboxylase is critical for non-small-cell lung cancer proliferation. The Journal of Clinical Investigation, 125(2), 687–698. https://doi.org/10.1172/JCI72873

Simsek, T., Kocabas, F., Zheng, J., DeBerardinis, R. J., Mahmoud, A. I., Olson, E. N., Schneider, J. W., Zhang, C. C., & Sadek, H. A. (2010). The Distinct Metabolic Profile of Hematopoietic Stem Cells Reflects Their Location in a Hypoxic Niche. Cell Stem Cell, 7(3), 380–390. https://doi.org/10.1016/j.stem.2010.07.011

Suda, T., Takubo, K., & Semenza, G. L. (2011). Metabolic regulation of hematopoietic stem cells in the hypoxic niche. Cell Stem Cell, 9(4), 298–310. https://doi.org/10.1016/j.stem.2011.09.010

Sun, J., Ramos, A., Chapman, B., Johnnidis, J. B., Le, L., Ho, Y.-J., Klein, A., Hofmann, O., & Camargo, F. D. (2014). Clonal dynamics of native haematopoiesis. Nature, 514(7522), 322–327. https://doi.org/10.1038/nature13824

Takizawa, H., Boettcher, S., & Manz, M. G. (2012). Demand-adapted regulation of early hematopoiesis in infection and inflammation. Blood, 119(13), 2991–3002. https://doi.org/10.1182/blood-2011-12-380113

Takubo, K., Nagamatsu, G., Kobayashi, C. I., Nakamura-Ishizu, A., Kobayashi, H., Ikeda, E., Goda, N., Rahimi, Y., Johnson, R. S., Soga, T., Hirao, A., Suematsu, M., & Suda, T. (2013). Regulation of glycolysis by Pdk functions as a metabolic checkpoint for cell cycle quiescence in hematopoietic stem cells. Cell Stem Cell, 12(1), 49–61. https://doi.org/10.1016/j.stem.2012.10.011

Tan, D. Q., & Suda, T. (2018). Reactive Oxygen Species and Mitochondrial Homeostasis as Regulators of Stem Cell Fate and Function. Antioxidants & Redox Signaling, 29(2), 149–168. https://doi.org/10.1089/ars.2017.7273

Thorvaldsdóttir, H., Robinson, J. T., & Mesirov, J. P. (2013). Integrative Genomics Viewer (IGV): High-performance genomics data visualization and exploration. Briefings in Bioinformatics, 14(2), 178–192. https://doi.org/10.1093/bib/bbs017

Tothova, Z., Kollipara, R., Huntly, B. J., Lee, B. H., Castrillon, D. H., Cullen, D. E., McDowell, E. P., Lazo-Kallanian, S., Williams, I. R., Sears, C., Armstrong, S. A., Passegué, E., DePinho, R. A., & Gilliland, D. G. (2007). FoxOs Are Critical Mediators of Hematopoietic Stem Cell Resistance to Physiologic Oxidative Stress. Cell, 128(2), 325–339. https://doi.org/10.1016/j.cell.2007.01.003

Trottier, M. D., Naaz, A., Li, Y., & Fraker, P. J. (2012). Enhancement of hematopoiesis and lymphopoiesis in diet-induced obese mice. Proceedings of the National Academy of Sciences, 109(20), 7622–7629. https://doi.org/10.1073/pnas.1205129109

Umemoto, T., Hashimoto, M., Matsumura, T., Nakamura-Ishizu, A., & Suda, T. (2018). Ca2+-mitochondria axis drives cell division in hematopoietic stem cells. The Journal of Experimental Medicine, 215(8), 2097–2113. https://doi.org/10.1084/jem.20180421

Vander Heiden, M. G., Cantley, L. C.,& Thompson, C. B. (2009). Understanding the Warburg effect: The metabolic requirements of cell proliferation. Science (New York, N.Y.), 324(5930), 1029–1033. https://doi.org/10.1126/science.1160809

Vanderperre, B., Herzig, S., Krznar, P., Hörl, M., Ammar, Z., Montessuit, S., Pierredon, S., Zamboni, N., & Martinou, J.-C. (2016). Embryonic Lethality of Mitochondrial Pyruvate Carrier 1 Deficient Mouse Can Be Rescued by a Ketogenic Diet. PLOS Genetics, 12(5), e1006056. https://doi.org/10.1371/journal.pgen.1006056

Vannini, N., Girotra, M., Naveiras, O., Nikitin, G., Campos, V., Giger, S., Roch, A., Auwerx, J., & Lutolf, M. P. (2016). Specification of haematopoietic stem cell fate via modulation of mitochondrial activity. Nature Communications, 7(1), 13125. https://doi.org/10.1038/ncomms13125

Ventura, A., Kirsch, D. G., McLaughlin, M. E., Tuveson, D. A., Grimm, J., Lintault, L., Newman, J., Reczek, E. E., Weissleder, R., & Jacks, T. (2007). Restoration of p53 function leads to tumour regression in vivo. Nature, 445(7128), 661–665. https://doi.org/10.1038/nature05541

Vigueira, P. A., McCommis, K. S., Schweitzer, G. G., Remedi, M. S., Chambers, K. T., Fu, X., McDonald, W. G., Cole, S. L., Colca, J. R., Kletzien, R. F., Burgess, S. C., & Finck, B. N. (2014). Mitochondrial pyruvate carrier 2 hypomorphism in mice leads to defects in glucose-stimulated insulin secretion. Cell Reports, 7(6), 2042–2053. https://doi.org/10.1016/j.celrep.2014.05.017

Wang, Y.-H., Israelsen, W. J., Lee, D., Yu, V. W. C., Jeanson, N. T., Clish, C. B., Cantley, L. C., Vander Heiden, M. G., & Scadden, D. T. (2014). Cell-State-Specific Metabolic Dependency in Hematopoiesis and Leukemogenesis. Cell, 158(6), 1309–1323. https://doi.org/10.1016/j.cell.2014.07.048

Warburg, O. (1925). The Metabolism of Carcinoma Cells. The Journal of Cancer Research, 9(1), 148–163. https://doi.org/10.1158/jcr.1925.148

Wright, D. E., Cheshier, S. H., Wagers, A. J., Randall, T. D., Christensen, J. L., & Weissman, I. L. (2001). Cyclophosphamide/granulocyte colony-stimulating factor causes selective mobilization of bone marrow hematopoietic stem cells into the blood after M phase of the cell cycle. Blood, 97(8), 2278–2285. https://doi.org/10.1182/blood.v97.8.2278

Yang, C., Ko, B., Hensley, C. T., Jiang, L., Wasti, A. T., Kim, J., Sudderth, J., Calvaruso, M. A., Lumata, L., Mitsche, M., Rutter, J., Merritt, M. E., & DeBerardinis, R. J. (2014). Glutamine Oxidation Maintains the TCA Cycle and Cell Survival during Impaired Mitochondrial Pyruvate Transport. Molecular Cell, 56(3), 414–424. https://doi.org/10.1016/j.molcel.2014.09.025

Yu, W., Liu, X., Shen, J., Jovanovic, O., Pohl, E. E., Gerson, S. L., Finkel, T., Broxmeyer, H. E., & Qu, C. (2012). Metabolic Regulation by the Mitochondrial Phosphatase PTPMT1 Is Required for Hematopoietic Stem Cell Differentiation. Stem Cell, 12(1), 62–74. https://doi.org/10.1016/j.stem.2012.11.022

